# High-frequency sorghum transformation toolkit enhances *Cas9* efficiency and expands promoter-editing capability with *SpRY*

**DOI:** 10.1101/2025.01.21.634149

**Authors:** Jianqiang Shen, Kiflom Aregawi, Sultana Anwar, Tamara Miller, Evan Groover, Mantra Rajkumar, David Savage, Peggy G. Lemaux

## Abstract

Sorghum bicolor L. (Moench), the fifth most important cereal crop internationally, is used as food, feed, forage, and fuels. Importantly, sorghum’s natural tolerance to environmental stresses leads to tolerance to climate variation. To optimize sorghum germplasm for crop improvement requires highly efficient genetic transformation and genome editing; however, sorghum has historically been recalcitrant to these genetic approaches. In this study, we report a high-efficiency engineering toolkit, optimized for genome editing via Agrobacterium-mediated transformation. Using CRISPR/Cas9-based editing machinery, the genetic tools led to editing efficiencies up to 95.7% when monocistronic guide RNAs targeted the phytoene desaturase (SbPDS) gene. We also tested a PAM (protospacer adjacent motif)-broadened Cas9 variant, intronized SpRY (ZmSpRYi), achievling comparable editing efficiencies. Using this toolkit permits exploitation of advanced editing tools, like prime editors, base editors, and targeted knock-in methods. These genetic advances offer new methods for sustainable crop improvement in sorghum and potentially other cereals.

**Technology readiness:** This study demonstrates the development of an integrated genome engineering platform that combines morphogenic gene-assisted transformation with CRISPR/Cas9- and SpRY-mediated mutagenesis for multiplex genome editing, establishing a Technology Readiness Level (TRL) of 4/5. This workflow enables the generation of allelic variants through editing of both functional and regulatory elements in the plant genome. Advancing toward TRL 5 will require additional validation in relevant sorghum genotypes, including elite varieties. Moreover, the PAM-flexible editor SpRY is still under optimization, particularly at non-canonical PAM sites, which may limit its application for the modification of certain cis-regulatory elements. Further improvements in precision genome editing, such as the integration of prime editing or targeted knock-in technologies, will also be necessary to enable desired and predictable modifications. With continued optimization and incorporation of these advanced tools, this robust genome engineering platform has the potential to accelerate plant breeding and enable basic research supporting plant-based solutions to address climate change.

## Introduction

Meeting future crop production demands, given more variable climate conditions and an increasing world population, requires higher crop yields with the same land usage. *Sorghum bicolor* L. (Monech) is an ideal crop for such future applications due to the diversity of its germplasm, particularly with respect to its multiple abiotic stress tolerances [1–3]. Breeding improved sorghum varieties, using precise genome editing, will accelerate development of improved cultivars [4–6]. However, genome editing with CRISPR/*Cas9* in sorghum has lagged in efficiency relative to other crops due to inefficient transformation systems for diverse genotypes, low editing percentage, and lack of optimized tools [7,8]. The current work seeks to address these stated limitations by utilizing a novel set of resources to achieve a high efficiency of sorghum transformation and genome editing.

Sorghum genome engineering has historically been a challenging process due to recalcitrance of most sorghum genotypes to *in vitro* tissue culture and a lack of technical expertise and effective tools for transformation [7]. A significant challenge for genome engineering in crop species is the requirement for germline transmission of mutations, necessitating gene delivery in pluripotent cells. Morphogenic factors, such as Babyboom (BBM) and Wuschel2 (WUS2) or WUS2 alone, increase the population of cells in target tissues that are competent for germline transmission [9–11]. Previously we demonstrated that transient expression from *Bbm* and *Wus2* during Agrobacterium-mediated gene delivery facilitated transformation of several recalcitrant sorghum varieties [8]. Concurrently, Che et al. demonstrated that during sorghum transformation incorporation of the *Wus2* gene alone increased the rate of CRISPR/*Cas9*-mediated knockout of *bmr,* the brown midrib gene [11]. The breakthrough in transformation technology due to morphogene-assisted transformation has greatly expanded the possibility of genotype-independent gene editing; however, developing tools for high efficiency genome editing with CRISPR remains a challenge.

The CRISPR/*Cas9* genome editing system, first discovered as a component of the adaptive immune response in *Streptococcus pyogenes* bacteria [12], is a powerful tool for genome editing in plants that has potential for functional genomics efforts and accelerated breeding [13]. However, editing in non-model plants lags in efficiency compared to animal and microbial systems, due to several factors, including the plant ‘regeneration bottleneck’ [7] and possibly sub-optimal activity of editors *in planta* [8,14]. Optimization of Cas endonuclease and guide RNA (gRNA) expression have been proposed to increase the efficiency of genome editing in sorghum [14]. In addition, use of Cas9 variants with expanded Protospacer Adjacent Motif (PAM) recognition sequences increases editing scopes [15–20]. To address these challenges further in Arabidopsis and tobacco, codon optimization of the Cas9 nuclease was used to increase gene knockout editing efficiency [21,22]. Use of specific promoters and gRNA transcriptional configurations was shown to increase sorghum editing efficiency, when the endogenous sorghum U6 promoter was used to drive gRNAs [23]. An additional limitation for introducing genetic variations at specific sites is the fact that the canonical *Cas9* endonuclease requires the NGG (N = A, T, C, G) PAM site, which limits the genome sites available for editing at sites with suitable PAM [24]. This limitation is especially relevant for engineering genomic regions such as *cis*-regulatory elements [25], where intra-populational sequence variations in promoter regions contribute significantly to phenotypic variations [26].

Several Cas9 variants have been engineered to overcome the PAM limitation of *SpCas9* [16,27,28]. *SpRY* endonuclease, an engineered variant of *Streptococcus pyogenes Cas9,* has a nearly PAM-less requirement and has been used for base editing and functional genetics in rice [19,20]. To demonstrate the utility of the SpRY endonuclease for editing cis-regulatory elements in sorghum, we tested its efficiency for editing the promoter of the *CYP79A1* gene, a step in dhurrin biosynthesis. Presence of dhurrin in sorghum limits its use as a forage because some varieties produce significant levels of this cyanogenic compound that can be fatal to livestock and accumulates under stress conditions [29–32]. Because dhurrin is also implicated in nitrogen storage and biotic stress tolerance [33,34], fine-tuning dhurrin biosynthesis, rather than fully eliminating it, is essential in balancing livestock safety with plant resilience and performance [30].

The present work addresses three main goals for advancing transformation and gene editing in *Sorghum bicolor*: increasing efficiency of CRISPR/*Cas9* editing, expanding available PAM recognition sites, and providing efficient genome editing capability. We developed a flexible functional genomics toolbox for Agrobacterium-mediated sorghum transformation and genome editing. The toolbox includes a binary T-DNA vector incorporating the *Wus2* gene, along with genes for *hygromycin B phosphotransferase* (*hph*) and *ZsGreen* (*ZsG*) used for selection and as a visual marker, respectively. For CRISPR/*Cas9* genome editing, we utilized an intronized Cas9 endonuclease in a shuttle vector, enabling easy insertion into the final binary vector of designed gRNA sequences and recombination. Additionally, for the first time in sorghum, we demonstrate editing with the nearly PAM-less Cas9 variant ZmSpRYi, employing both poly- and monocistronic gRNA configurations to target edits in the promoter region of the *CYP79A1* dhurrin biosynthesis gene. Overall, this high-efficiency toolbox increases the scalability of genome editing in sorghum, enhancing future efforts for accelerated breeding and synthetic biology.

## Results

### Design of high efficiency binary and shuttle vectors

Our two-step strategy for vector development involves gRNA validation in protoplasts, followed by stable plant genetic transformation (**Figure 1A**). This advance in using protoplasts can enhance efficiency and accessibility for researchers to perform successful sorghum transformation and editing. The editing machinery is integrated in shuttle vectors, which enable gRNA cloning with insertion of either a poly- or monocistronic configuration (**Figure 1B**). The gRNA scaffold and Pol III promoters necessary for driving gRNA expression are supplied by the Level 0 constructs (**Figure S1**). The Level 0 and shuttle constructs are sequentially assembled to form complete gRNA configurations, which are transfected into sorghum protoplasts to evaluate editing efficiency of the gRNAs [35,36].

**Figure 1.**
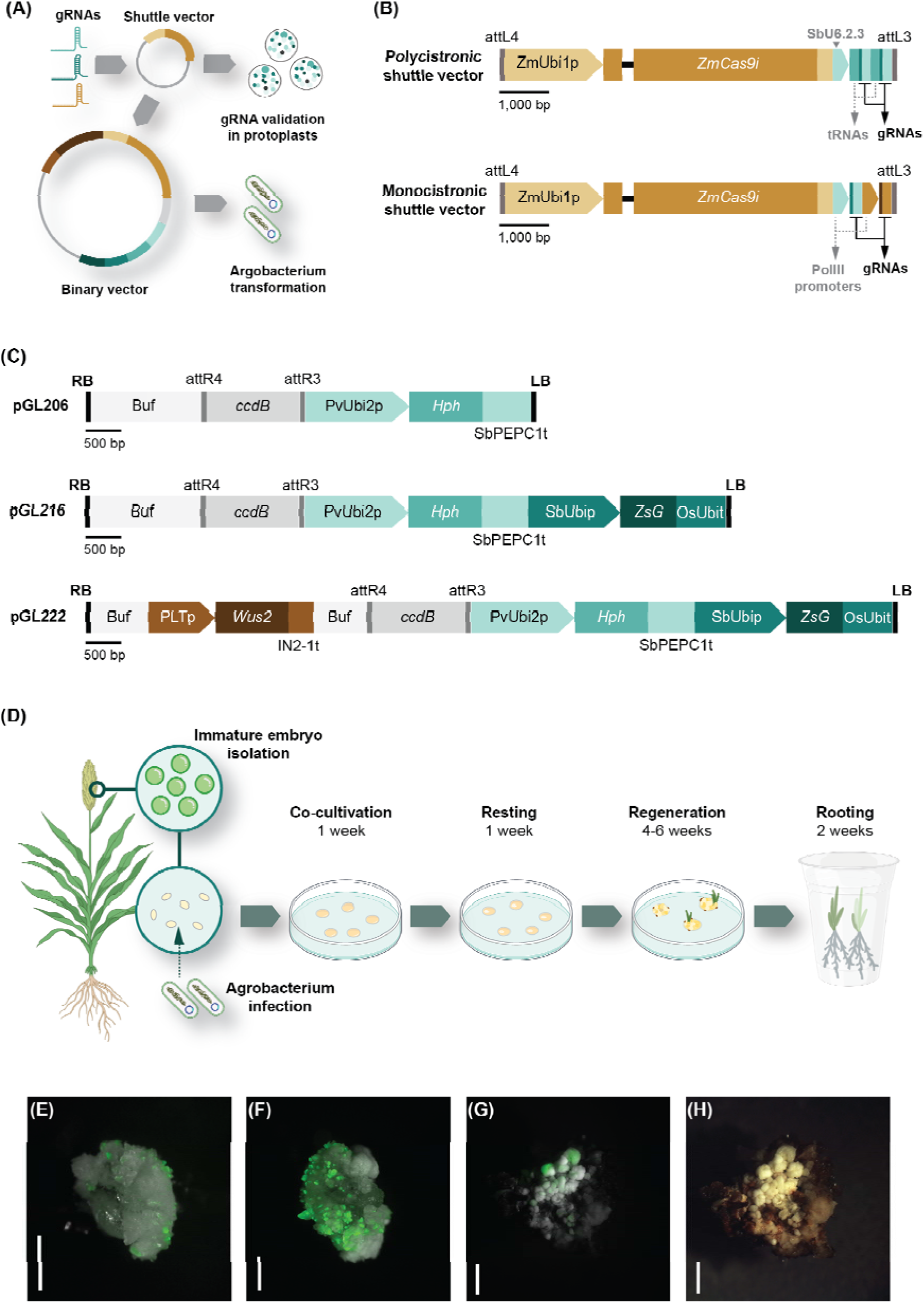
Workflow for construct design used for sorghum immature embryo transformation. **(A)** Diagram illustrating the workflow for construct creation and usage. **(B)** Maps of shuttle vectors. Polycistronic shuttle vector contains a polycistronic gRNA transcription unit with tRNA spacers; monocistronic shuttle vector contains monocistronic gRNA transcription units driven by Pol III promoters. Rectangles with arrows indicate promoters. Other rectangles indicate exons (dark-colored) or terminators (light-colored). Black horizontal lines indicate introns. Grey vertical lines indicate Gateway recombination sites. **(C)** Maps of binary vectors. pGL206 represents the basic binary vector backbone. pGL216 is designed for testing the function of morphogenic genes in enhancing transformation frequencies. pGL222 is a binary vector backbone used in genome editing applications. Rectangles with arrows indicate promoters. Solid rectangles indicate exons (dark-colored) or terminators (light-colored). Light gray rectangles denote buffer regions; dark gray rectangles represent *ccdB* genes. Black vertical lines indicate T-DNA borders; gray vertical lines indicate Gateway recombination sites; black vertical lines indicate left and right borders. *Hygromycin phosphotransferase* (*Hph*) gene is driven by switchgrass *ubiquitin 2* promoter (*PvUbi2*p) and terminated by sorghum *phosphoenolpyruvate carboxylase* terminator (*SbPEPC1*t); *ZsGreen1* (*ZsG*) gene is driven by sorghum ubiquitin promoter (*SbUbi*p) and terminated by rice *ubiquitin* terminator (*OsUbi*t); Maize *Wuschel2* (*Wus2*) gene is driven by maize *phospholipid protein* promoter (*PLT*p) and terminated by maize *IN2-1* terminator (*IN2-1*t). **(D)** Diagram illustrating the transformation workflow, from immature embryo isolation through rooted regenerants. **(E-H)** Sorghum tissues at various stages post-Agrobacterium infection. **(E)** Immature embryos one week after infection; **(F)** two weeks after infection and **(G)** four weeks after infection, visualized with overlays of brightfield and fluorescence images captured with a Zeiss AxioZoom V16 low-magnification epifluorescence microscope. Scale bars indicate 2 mm. **(H)** Sorghum tissue four weeks after Agrobacterium infection, with two weeks of exposure to hygromycin selection. Scale bar indicates 2 mm.

To enhance efficiency of sorghum transformation, a series of binary constructs were developed, all of which are compatible with Gateway LR recombination and tailored to allow incorporation of genes of interest for multiple applications (**Figure 1C**). A hygromycin resistance gene (*hph*), used as a selectable marker in the foundational construct, pGL206 (**Figure 1C**), was chosen because of the stringent selection properties of hygromycin during *in vitro* tissue culture of sorghum [37]. pGL206 was used as a construct for testing a candidate morphogenic gene, i.e., maize *Wuschel2* (*Wus2*). To facilitate tracking of somatic embryogenesis and plantlet development, a visible marker gene *ZsGreen1,* driven by the sorghum ubiquitin promoter, was incorporated near the T-DNA left border in pGL216. Building on previous findings of enhanced editing efficiency achieved with integration of maize *Wus2* [11], pGL216 was assembled with *Wus2*, driven by the maize phospholipid transfer protein promoter (pLTP). This resulted in pGL222 (**Figure 1C**), a binary vector optimized for delivering genome-editing machinery. Expression of ZsGreen validates presence of at least one complete T-DNA copy and permits easy visualization of T-DNA segregation in T_1_ generations. The constructs developed provide a robust framework for improving transformation and genome-editing efficiencies in sorghum.

### High transformation frequency achieved through *Wus2* integration

To evaluate transformation frequency in genome editing trials, we knocked out the *Sorghum bicolor phytoene desaturase* (*SbPDS*) gene, resulting in easy visualization of an albino phenotype [8]. Previous studies indicated editing efficiency may vary in wheat (*Triticum aestivum*) and setaria (*Setaria viridis*) depending on the nature of the configurations of gRNAs in C3 and C4 grasses, respectively [38,39]; however, this has not been thoroughly investigated in sorghum using CRISPR/*Cas9*. To optimize editing efficiency, constructs were designed containing two gRNAs under two transcriptional configurations, polycistronic and monocistronic (**Figure 1B**). In the polycistronic construct, gRNAs were driven by the sorghum snoRNA U6.2.3 promoter [23]. In the monocistronic configuration, gRNAs were expressed using the snoRNA U6.2.3 and snoRNA U6.3.1 promoters [23]. Both validated gRNA transcriptional configurations were combined with a Cas9 expression cassette and integrated into the binary vector pGL222, resulting in two editing constructs, pGL220 (polycistronic) and pGL221 (monocistronic) (**Figure S2**). The constructs, containing the morphogenic gene, *Wuschel2* (*Wus2*), and the selectable marker gene, *hph*, were used to assess the impact on editing efficiency of the two gRNA transcriptional configurations.

Both pGL220 and 221 were transformed into sorghum immature embryos (IEs) using a modified protocol [8]. Following Agrobacterium infection, IEs were maintained at 25°C for one week on co-cultivation medium. In the current approach (**Figure 1D**), unlike *Bbm*/*Wus2* assisted transformation, the *Wus2* expression cassette does not need to be excised because *Wus2* integration does not lead to severely abnormal phenotypes. Previous studies in wheat indicated that using higher temperatures at this stage enhanced mutagenesis efficiency when employing CRISPR/*Cas9* [40]. To explore this in sorghum, tissues were incubated, starting at the resting stage, at either 25°C or 28°C until plant regeneration began. Development of IEs was monitored to evaluate responses to *Wus2* expression at 1-, 2-, and 4-week intervals post-infection (**Figures 1E–H**). Fluorescence was detected as early as three days after infection (**Figure 1E**) and peaked at the end of the resting stage (**Figure 1F**). Somatic embryogenesis was noticeable at two weeks, with regenerated plantlets forming by week four (**Figures 1G-H**). Prior to transferring to rooting media, fluorescence microscopy was used to screen for ZsGreen expression, confirming integration of full-length T-DNA, given that the ZsGreen expression cassette is located adjacent to the left border.

Putative transgenic plantlets were analyzed via PCR to further confirm T-DNA integration and determine transformation frequencies. Transformation frequencies and editing efficiencies were calculated using two strategies. The first was to calculate frequency using only one transgenic plant regenerated from each IE; the second used all regenerated transgenic plants from each IE. Both approaches have advantages. Considering a single plant per IE is helpful when using an overexpression construct in order to get independent transgenic events. Conversely, calculating transformation frequency by considering all regenerated plantlets from each IE, as in the current approach, is beneficial when performing genome editing because multiple independently edited events can be regenerated from a single IE and may have unique edits.

For the monocistronic strategy at 28°C, when considering only one plant per IE, we generated 50 transgenic plants from two replications, obtaining a transformation frequency of 24.4% (50/205) (**Table 1)**. For the polycistronic strategy an average transformation frequency was 26.9% (81/301). It is important to note that issues incubated at 28°C appeared stressed and showed more necrosis than those at 25°C (**Figure S3**). Because of these phenotypic differences, more plants were regenerated from tissues kept at 25°C, increasing transformation frequency for the monocistronic strategy to 31.1% versus 24.4% for tissues kept at 28°C. When all regenerated transgenic plants were considered, transformation frequency substantially increased (**Table 2**). At 25°C, the monocistronic strategy yielded transformation frequency of 79.4% (143/180). Additionally, at 28°C, it led to notable differences in transformation frequency, 66.8% (137/205) (**Table 2**). This decrease in frequency may be due to the fact that tissues kept at 28°C showed more necrosis than those kept at 25°C (**Figure S3**), thereby reducing the number of regenerated plants for the monocistronic strategy thus lowering transformation frequencies (**Table 1**).

**Table 1.** Transformation, editing and knockout efficiencies of Wus2-assisted transformation, based on one plantlet per IE.

| gRNA system (Construct) | Temp (°C) | Replication | # IEs used | # Regenerants | # Escapes (%)<sup>†</sup> | + for hph<sup>‡</sup> | Transformation frequency (%)<sup>§</sup> | # Edited plants | Editing efficiency (%)<sup>¶</sup> | # Plants with knockout<sup>††</sup> | Knockout efficiency (%)<sup>‡‡</sup> |
| --- | --- | --- | --- | --- | --- | --- | --- | --- | --- | --- | --- |
| Polycistronic (pGL220) | 28 | 1st | 132 | 47 | 6 (12.8) | 41 | **31.0** | 24 | **58.5** | 24 | **58.5** |
|  |  | 2nd | 169 | 44 | 4 (9.1) | 40 | **23.7** | 25 | **62.5** | 25 | **62.5** |
|  |  | Total | 301 | 91 | 10(10.9) | 81 | **26.9 (%)** | 49 | **60.5 (%)** | 49 | **60.5 (%)** |
|  | 25 | 1st | 143 | 30 | 4 (13.3) | 26 | **18.2** | 17 | **65.4** | 17 | **65.4** |
| Monocistronic (pGL221) | 28 | 1st | 93 | 24 | 1 (4.2) | 23 | **24.7** | 22 | **95.7** | 22 | **95.7** |
|  |  | 2nd | 112 | 28 | 1 (3.6) | 27 | **24.1** | 26 | **96.3** | 26 | **96.3** |
|  |  | Total | 205 | 52 | 2(3.8) | 50 | **24.4 (%)** | 48 | **96.0 (%)** | 48 | **96.0 (%)** |
|  | 25 | 1st | 105 | 34 | 1 (2.9) | 33 | **31.4** | 33 | **100.0** | 33 | **100.0** |
|  |  | 2nd | 75 | 23 | 0(0) | 23 | **30.7** | 21 | **91.3** | 21 | **91.3** |
|  |  | Total | 180 | 57 | 1(1.8) | 56 | **31.1 (%)** | 54 | **96.4 (%)** | 54 | **96.4 (%)** |

<sup>†</sup> # Escapes (%) = # and % of regenerated plants without transgene (T-DNA) = (# Escapes/# Regenerants) 100
<sup>‡</sup> + = # plants PCR-positive for *hph*
<sup>§</sup> Transformation frequency (%) = (# + for *hph* divided by # IEs used) x 100
<sup>¶</sup> Editing efficiency (%) = (# edited plants divided by # plants + for *hph*) x 100
<sup>††</sup> # Plants with knockout = # plants with frameshift mutation and albino phenotype
<sup>‡‡</sup> Knockout efficiency(%) = (# plants with knockout divided by # + for *hph*) x 100

**Table 2.** Transformation, editing and knockout efficiencies of Wus2-assisted transformation, based on multiple regenerated plantlets from each IE.

| gRNA system (Construct) | Temp (°C). | Replications | # IEs used | # Regenerants | # Escapes (%) <sup>†</sup> | + for *hph* <sup>‡</sup> | Transformation frequency (%) <sup>§</sup> | # Edited plants | Editing efficiency (%) <sup>¶</sup> | # Plants with Knockout <sup>††</sup> | Knockout efficiency (%) <sup>‡‡</sup> |
| --- | --- | --- | --- | --- | --- | --- | --- | --- | --- | --- | --- |
| Polycistronic (pGL220) | 28 | 1st | 132 | 60 | 6 (10) | 54 | **40.9** | 30 | **55.6** | 30 | **55.6** |
|  |  | 2nd | 169 | 48 | 4 (8.3) | 44 | **26.0** | 27 | **61.4** | 27 | **61.4** |
|  |  | Total | 301 | 108 | 10(9.2) | 98 | **32.6** | 57 | **58.2** | 57 | **58.2** |
|  | 25 | 1st | 143 | 31 | 4 (12.9) | 27 | **18.9 (%)** | 17 | **63.0 (%)** | 17 | **63.0** |
|  |  | 2nd |  |  |  |  |  |  |  |  |  |
|  |  | Total |  |  |  |  |  |  |  |  |  |
| Monocistronic (pGL221) | 28 | 1st | 93 | 64 | 1 (1.6) | 63 | **67.7** | 61 | **96.8** | 60 | **95.2** |
|  |  | 2nd | 112 | 77 | 3 (3.9) | 74 | **66.1** | 66 | **89.2** | 63 | **85.1** |
|  |  | Total | 205 | 141 | 4(2.8) | 137 | **66.8 (%)** | 127 | **92.7 (%)** | 123 | **89.8** |
|  | 25 | 1st | 105 | 93 | 6 (6.5) | 87 | **82.9** | 87 | **100.0** | 84 | **96.6** |
|  |  | 2nd | 75 | 60 | 4(6.7 ) | 56 | **74.7** | 51 | **91.1** | 49 | **87.5** |
|  |  | Total | 180 | 153 | 10(6.5) | 143 | **79.4 (%)** | 138 | **96.5 (%)** | 133 | **93.0** |

<sup>†</sup> # Escapes (%) = # and % of regenerated plants without transgene (T-DNA) = (# Escapes/# Regenerants) X 100
<sup>‡</sup> + = # Plants PCR-positive for *hph*
<sup>§</sup> Transformation frequency (%) = (# + for *hph* divided by # IEs used) x 100
<sup>¶</sup> Editing efficiency (%) = (# Edited plants divided by # plants + for *hph*) x 100
<sup>††</sup> # plants with knockout = # plants with frameshift mutation and albino phenotype
<sup>‡‡</sup> Knockout efficiency (%) = (# Plants with knockout divided by # + for *hph*) x 100

### Enhancing CRISPR/Cas9 editing efficiency using WUS2

Previous work demonstrated that various factors can be used to improve CRISPR/*Cas9* editing efficiency. One factor is intronized and codon-optimized *Cas9* [21,22]. Using this *Cas9* to improve genome editing might result from enhanced mRNA stability and translation [22]. Another possible factor is the nature of promoters driving gRNAs (**Figure 1B, 2A**), as they play an important role in regulating expression of gRNAs [23,41]. Additionally incubating tissues at higher temperatures might lead to improved editing efficiency [40,42]. Using a *Wus2*-integrated transformation strategy resulted in improved CRISPR/*Cas9* gene knockout frequency, which might be due to enhanced cell division and/or chromatin accessibility [11]. In our study, different factors were used that could lead to improved genome editing and knockout efficiencies. The approaches include using: (i) the *Wus2*-integrated transformation strategy, (ii) a codon-optimized and intronized *Cas9*, (iii) monocistronic and/or polycistronic tRNA-gRNA expression cassettes, (iv) the *SbU6* 2.3 and *SbU6* 3.1 promoters driving two gRNAs, and/or (v) two temperatures, 25°C and 28°C, during certain tissue culture steps.

To assess the impact of the various factors stated above, we focused on *SbPDS* due to the ability to observe varied albino phenotypes **(Figure 2B).** This phenotype was used to visually identify plants with *SbPDS* gene knockouts and to calculate knockout efficiencies. In addition, we amplified the *SbPDS* gene fragments from transgenic plants and used Sanger sequencing to investigate genome editing outcomes. Results were further analyzed using the Inference of <u>C</u>RISPR <u>E</u>dits (ICE) Analysis Tool [43]. Editing outcomes for individual plants, as measured by knockout scores from ICE, ranged from 0% to 100% (**Figure 2C**). Samples with knockout scores greater than 5% were considered successfully edited. Genotypic editing efficiency (**Tables 1, 2**) was calculated as the percentage of edited plants among all transformants for that condition, hereafter this will be termed editing efficiency. Phenotypic knockout efficiency (**Tables 1, 2**) was defined as the percentage of plants showing an albino phenotype among all transformants, hereafter this will be termed knockout efficiency. The combined effects of the factors mentioned in the preceding paragraph resulted in substantially improved average editing and knockout efficiencies reaching 65.4% (17/26) for polycistronic and 96.4% (54/56) for monocistronic configurations in tissues maintained at 25°C (**Table1**), compared to efficiencies of 22.6% and 16.7%, respectively, in our previous report [8]. Incubation at 28°C versus 25°C did not yield notable differences in editing and knockout efficiencies (**Table 1**). Interestingly, somatic embryos cultured at 25°C appeared less stressed under hygromycin selection, perhaps due to reduced selection pressure relative to those at 28°C, as indicated by less tissue browning (**Figure S3**). Furthermore, analysis of knockout scores for individual gRNAs revealed that gRNA1.1 in the polycistronic constructs exhibited a notably reduced knockout score (**Figure 2C**), possibly due to suboptimal processing efficiency during tRNA-mediated autocleavage. Overall, our results highlight the robustness of the monocistronic strategy, although the polycistronic design can also deliver reasonably high editing efficiencies.

**Figure 2.**
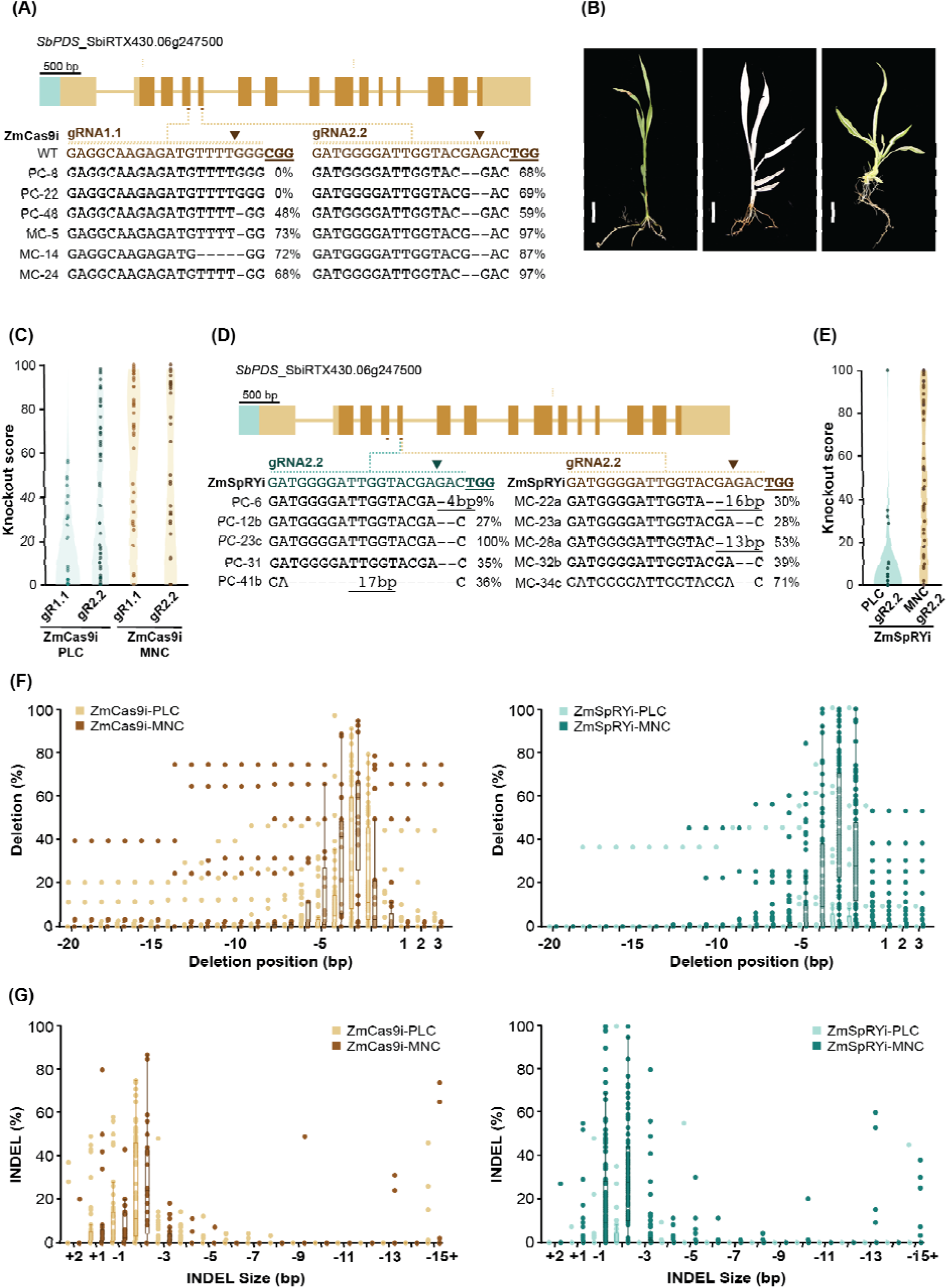
CRISPR/*Cas9* and CRISPR/*SpRY*-mediated editing of the *SbPDS* gene using *Wus2*-integrated transformation. **(A)** Top panel: schematic representation of the structure of *SbPDS*, depicting *SbPDS* exons and gRNA target sequences. Light green rectangle indicates promoter. Light yellow rectangles represent untranslated regions. Dark gold rectangles represent exons. Gold lines indicate introns. Guide RNAs are indicated by brown lines. Bottom panel: representative genotypes of PDS edits in T_0_ generation. Underlined sequences indicate gRNA Protospacer Adjacent Motif (PAM) sequences, and triangles represent potential double-strand break sites. Knockout scores for each gRNA are shown after each editing outcome. Knockout scores in percentages represent the dominant genotypes identified from amplicon sequencing, followed by analysis using Synthego Inference of CRISPR Edits (ICE) tools. **(B)** Phenotypes of PDS gene-edited plants. The left panel shows a non-edited plantlet; middle panel shows a fully albino plantlet; right panel shows a chimeric albino plantlet. Images were taken four weeks after initiation of rooting. Scale bar = 2 cm. **(C)** Violin plot showing the distribution of ZmCas9i-mediated knockout scores obtained from Sanger sequencing followed by Synthego ICE analysis. For PDS editing with polycistronic gRNA configurations (PLC), n=76; for PDS editing with monocistronic configurations (MNC), n=41. **(D)** Representative genotypes on gRNA2.2, resulting from ZmSpRYi-mediated editing of the SbPDS gene in the T_0_ generation. Underlined sequences indicate gRNA PAM sequences; triangles represent potential double-strand break sites. Knockout scores for each gRNA are shown after each editing outcome. Knockout scores in percentage represent the dominant genotypes identified from amplicon sequencing, followed by analysis using Synthego ICE tools. For PDS editing with the polycistronic gRNA configuration, results are presented in the left panel; for PDS editing with monocistronic gRNA configuration, results are shown in the right panel. **(E)** Violin plot showing distribution of ZmSpRYi-mediated knockout scores for *PDS* gRNA2.2 edits mediated by CRISPR/SpRY, comparing polycistronic and monocistronic gRNA expression cassettes. For PLC, n = 68; for MNC, n = 92. **(F)** Comparison of deletion positions between ZmCas9i and ZmSpRYi at gRNA2.2 (NGG PAM sites) in T_0_ plantlets. Each dot represents a biological replicate. Distribution of deletion percentage from gRNA PLC and MNC configurations are shown by highlighting the median and quartiles of each. ZmCas9i-PLC, n=76; ZmCas9i-MNC, n=41. ZmSpRYi-PLC, n = 68; ZmSpRYi-MNC, n = 92. **(G)** Comparison of indel size between ZmCas9i and ZmSpRYi at gRNA2.2 (NGG PAM sites) in T_0_ plantlets. Each dot represents a biological replicate. Distribution of indel percentages from gRNA PLC and MNC configurations are shown by highlighting the median and quartiles of them. ZmCas9i-PLC, n=76; ZmCas9i-MNC, n=41. ZmSpRYi-PLC, n = 68; ZmSpRYi-MNC, n = 92.

T_0_ plants were phenotyped based on the protortion of albino sectors observed. Fully albino plantlets were scored as 100%, whereas full green plants as 0% represented. These phenotypic assessments were compared with the knockout scores gathered through the ICE analysis, enabling effective identification and quantification of the degree of penetrance and frequencies of mutations in plantlets. For most plants, phenotypic and genotypic scores were highly consistent, with a few discrepancies, such as in pGL220-2, -20 & -34 (**Table S2**). Overall, the two metrics showed a strong positive correlation, with a Pearson’s correlation coefficient of 0.81.

This correlation underscores the value of the ICE analysis as a reliable and affordable tool for estimating penetrance based on genotypic data. particularly in cases where gene knockouts do not produce visually detectable phenotypes in the T_0_ generation.

### High-efficiency editing using PAM-less Cas9 variant, ZmSpRYi

Previous studies demonstrated that CRISPR/*Cas9*-mediated editing of promoters can generate diverse *cis*-regulatory alleles, providing valuable quantitative variation for crop breeding applications [44–46]. However, the standard CRISPR/*Cas9* system relies on canonical gRNAs with NGG as PAM sequences, which are less likely in promoter or 5’ untranslated regions. In addition, CRISPR/*Cas9*-mediated edits in our study predominantly resulted in insertions or deletions of less than 3 bp (**Figure 2A**). These small modifications also present a challenge when targeting *cis*-regulatory elements, which can span long stretches of DNA. These regions can include multiple coordinated DNA motifs, which may not be effectively disrupted by such small edits [47]. Taken together, these observations indicate a limitation of CRISPR/Cas9 in editing cis-regulatory modules.

To expand the genome editing scope and to identify options with a less constrained PAM sequence, a range of nucleases were considered, including *Cas9* variants and *Cas9* orthologs [16–20]. We focused on a PAM-flexible *Cas9* variant, *SpRY* [24], which enables more frequent cis-regulatory element editing and expands genome engineering in AT-rich regions. To create a maize codon-optimized *SpRY* variant, *ZmSpRYi,* targeted mutations were introduced in the intronized *ZmCas9i* by site-directed mutagenesis (**Figure S4**). Previous studies suggested that *SpRY* is a less efficient endonuclease due to modified DNA unwinding activity [48]. To evaluate the editing efficiency of ZmSpRYi, we did a direct comparison between ZmSpRYi and ZmCas9i, using the same gRNAs for both (**Figure 2A, D**). Two constructs, featuring polycistronic and monocistronic configurations of gRNAs, were transformed into sorghum. To assess editing efficiency of ZmSpRYi, all independent transformants were genotyped by PCR amplification, followed by Sanger sequencing. Analysis of the sequencing data using ICE revealed that, when using the polycistronic transcription system, 26.7% (16/60) of independent transformants were edited whereas, strikingly when using the monocistronic system, 88.0% (81/ 92) of transformants were edited (**Table 3**, **Figure 2D**). Interestingly, the gRNA1.1 failed to induce any edits, suggesting that ZmSpRYi may exhibit distinct editing activity or property, compared to ZmCas9i. In contrast, the knockout score distribution of gRNA2.2 in the monocistronic configuration showed a robust editing efficiency (85.9%) versus the polycistronic configuration (26.7%) (**Table 3**), indicating that gRNAs expressed from monocistronic configurations are more efficient This is consistent with the findings when using CRISPR/Cas9-mediated genome editing (**Table 1**, **Figure 2C**). To investigate the editing characteristics of ZmSpRYi, we compared the editing profile of ZmCas9i and ZmSpRYi using amplicon sequencing from stable edits. Analysis of editing outcomes for gRNA2.2 revealed that deletions peaked at the third nucleotide upstream of the NGG site when both ZmCas9i and ZmSpRYi were used (**Figure 2F**). ZmSpRYi, however, generated a larger deletion range, spanning from -8 to +3, particularly with the gRNA monocistronic configuration (**Figure 2F**). Further, examination of indel sizes showed that both nucleases predominantly produced 2-bp deletions (**Figure 2G**), although deletions mediated by ZmSpRYi were slightly larger than those from ZmCas9i. These findings highlight the enhanced editing efficiency of the monocistronic system in sorghum when using either ZmCas9i or ZmSpRYi. Moreover, ZmSpRYi directed by NGG gRNAs may induce a wider spectrum of deletions, making it a promising tool for disrupting cis-regulatory elements.

**Table 3.** Transformation, editing and knockout efficiencies of *SpRY* using *Wus2*-assisted transformation.

| gRNA system<br>(Construct) | Replications | # IEs<br>used | # Regenerants | # Escapes<br>(%)<sup>†</sup> | + for<br>*hph*<sup>‡</sup> | Transformation<br>frequency (%)<sup>§</sup> | # Edited<br>plants | Editing<br>efficiency (%)<sup>¶</sup> | # Plants with<br>Knockout<sup>††</sup> | Knockout<br>efficiency (%) |
| --- | --- | --- | --- | --- | --- | --- | --- | --- | --- | --- |
| Polycistronic<br>(pGL226) | 1<sup>st</sup> | 107 | 46 | 3 (6.5) | 43 | 40.2 | 11 | 25.6 | 11 | 25.6 |
|  | 2<sup>nd</sup> | 60 | 21 | 4 (19.0) | 17 | 28.3 | 5 | 29.4 | 5 | 29.4 |
| Monocistronic<br>(pGL227) | 1<sup>st</sup> | 119 | 49 | 0 (0) | 49 | 41.2 | 40 | 81.6 | 40 | 81.6 |
|  | 2<sup>nd</sup> | 82 | 46 | 3 (6.5) | 43 | 52.4 | 41 | 95.3 | 39 | 90.7 |

<sup>†</sup> # Escapes (%) = # and % of regenerated plants without transgene (T-DNA) = (# Escapes/# Regenerants) X 100
<sup>‡</sup> + = # Plants PCR-positive for *hph*
<sup>§</sup> Transformation frequency (%) = (# + for *hph* divided by # IEs used) x 100
<sup>¶</sup> Editing efficiency (%) = (# Edited plants divided by # plants + for *hph*) x 100
<sup>††</sup> # Plants with knockout = # Plants with frameshift mutations and albino phenotypes
<sup>‡‡</sup> Knockout efficiency (%) = (# Plants with knockout divided by # plants + for *hph*) x 100

### Editing PDS with ZmSpRYi using non-canonical gRNAs

Canonical gRNAs for Cas9-mediated genome editing require NGG (N = A, T, C, G) as a PAM sequence, whereas non-canonical PAMs refer to all other sequences that can be recognized and bound by SpRY. To assess activity of gRNAs with noncanonical PAMs in combination with ZmSpRYi within AT-rich regions of the *PDS* promoter and exons, we designed six gRNAs with NAA, NAT, NTA or NTT PAMs (**Figure 3A**). Three gRNAs were positioned 168 bp, 100 bp, and 66 bp upstream of the transcription start site, aiming to disrupt translational regulation. The remaining three targeted coding sequences, two in the first exon and one in the second exon, were positioned to interfere with protein translation (**Figure 3A**). To optimize gRNA expression in monocots, we employed a variety of U6 and U3 snoRNA promoters derived from various grass species (**Figure S1**). All editing components, including ZmSpRYi and the six gRNAs, were delivered via Agrobacterium-mediated transformation.

**Figure 3.**
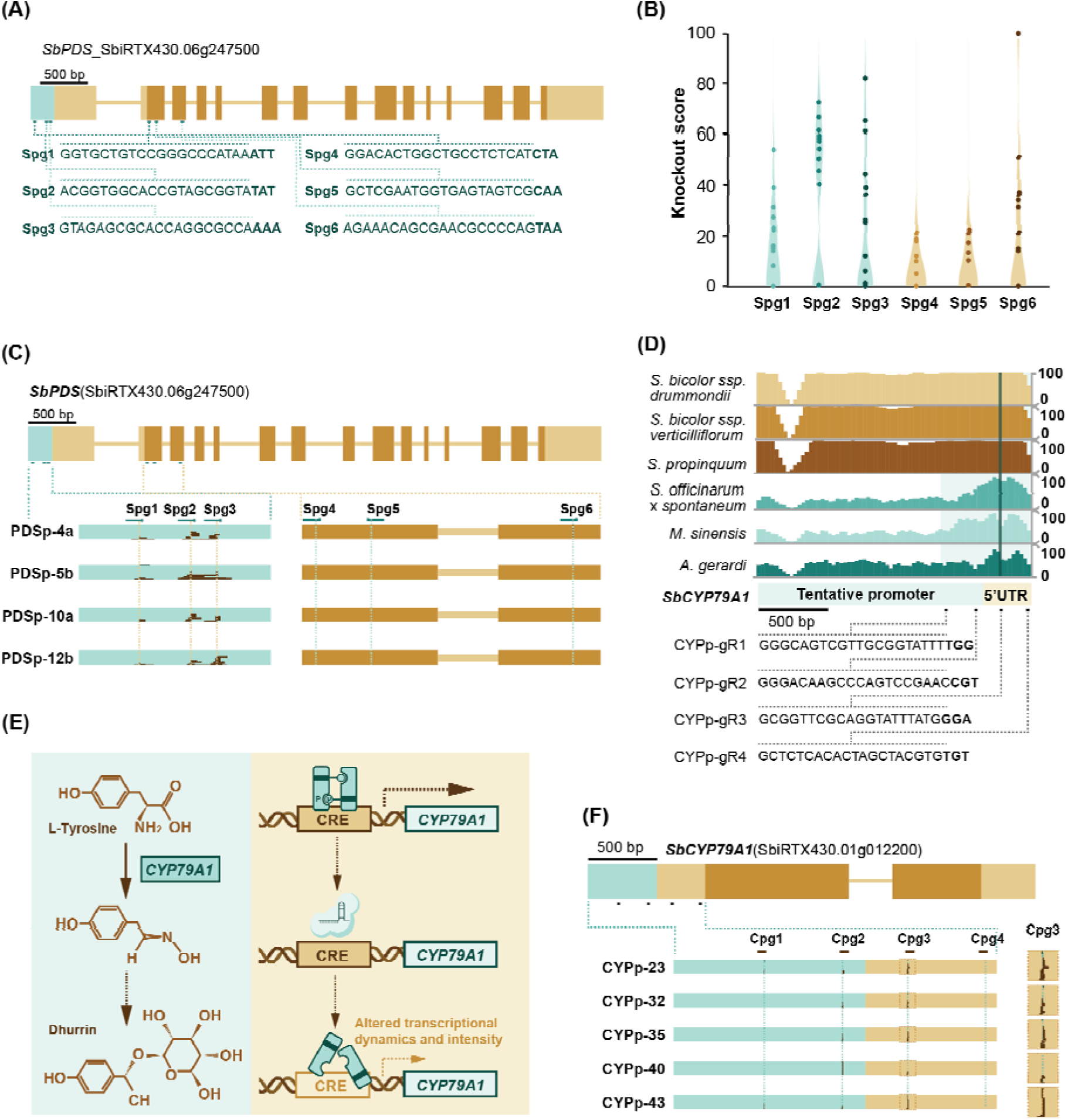
CRISPR/*SpRY*-mediated editing of cis-regulatory elements in promoter region. **(A)** Schematic representation of CRISPR/*SpRY* target sites on the *SbPDS* gene. A light green rectangle represents the promoter region; light gold rectangles indicate untranslated regions (UTRs); dark gold rectangles denote exons; connecting light gold lines represent introns. Guide RNA (gRNA) target sites are shown as green lines, with PAM sequences highlighted in bold. *SbPDS* gRNA (Spg) indicates the gRNA used for editing. **(B)** Violin plot showing distribution of individual knockout scores, represented as dots, for non-canonical gRNAs targeting *SbPDS*, as mediated by CRISPR/*SpRY*. *n* = 31. Spg was defined in the figure 3A legend. **(C)** Genotypic outcomes of zSpRYi-mediated editing targeting of the *SbPDS* promoter and coding sequences. *Top panel:* Diagram of *SbPDS* gene structure. A light green rectangle represents the promoter region; light gold rectangles indicate untranslated regions (UTRs); dark gold rectangles denote exons; connecting light gold lines represent introns. gRNA target sites are indicated by green dots. *Bottom panels:* Representative T genotypes and mutation patterns. *Left:* Genotypes from editing cis-regulatory elements in the promoter region. *Right:* Genotypes from edits within the exons. Vertical dashed lines indicate predicted double-strand break sites. Black rectangles represent deletions introduced via DNA repair; the width corresponds to indel length, and the height indicates the frequency of edited alleles, based on Synthego ICE analysis. Spg: *<u>S</u>b<u>P</u>DS* <u>g</u>RNA. **(D)** Conservation of *CYP79A1* promoter regions across C_4_ grasses. Bar graph shows sequence similarity between the *SbCYP79A1* promoter (from *Sorghum bicolor* genotype RTx430) and orthologous regions from *Miscanthus sinensis*, *Saccharum officinarum*, and *Andropogon gerardii*. Promoter regions analyzed span 2,000 bp upstream of the translation start site, with each bar representing a 30-nt window. gRNA locations within the promoter and UTR are indicated by black dots; PAM sequences are in bold. **(E)** Overview of the dhurrin biosynthetic pathway and cis-regulatory element editing strategy. *Left panel:* Simplified pathway highlighting *CYP79A1* as the initial step in dhurrin biosynthesis. *Right panel:* Workflow illustrating the use of cis-regulatory element editing to attempt to reduce dhurrin content by targeting the *SbCYP79A1* promoter and UTR. **(F)** Genotypic outcomes of zSpRYi-mediated editing of the *SbCYP79A1* promoter and UTR. *Top panel:* Gene diagram showing promoter, UTR, exons, and gRNA target sites (dots below the gene structure). A light green rectangle represents the promoter region; light gold rectangles indicate untranslated regions (UTRs); dark gold rectangles denote exons; connecting light gold lines represent introns. gRNA target sites are indicated by green dots. *Bottom panels:* Representative T genotypes reflecting mutation patterns from cis-regulatory region editing. Deletions are marked by dark brown rectangles. The width of the rectangles corresponds to indel length, and the height indicates the frequency of edited alleles, based on Synthego ICE analysis. A magnified view of editing outcomes from the cpg3 gRNA is also shown. Cpg: <u>C</u>YP79A1 <u>p</u>romoter <u>g</u>RNA.

A previous study reported that the reduced PAM specificity of SpRY might compromise editing efficiency, potentially due to decreased gRNA-target binding stability and inefficient R-loop formation [49]. To evaluate the *in vivo* activity of ZmSpRYi, the T_0_ regenerants described above were genotyped. Of the 64 independent transformants analyzed, 31 exhibited mutations at one or more gRNAs target sites, resulting in an overall editing efficiency of 48.4% (**Table 4**). Notably, ZmSpRYi-induced edits frequently resulted in larger deletions within the promoter region (**Figure S5**), reflecting a distinct mutational profile compared with ZmCas9i, which generally favors shorter indels in sorghum. Interestingly, Spg2 and Spg3, separated by 44 bp, exhibited markedly higher dropout frequencies (**Figure S5**). The tendency of ZmSpRYi to generate larger deletions may be particularly advantageous for promoter editing, as complete disruption of *cis*-regulatory elements often requires near-total removal to achieve functional inactivation [50]. Furthermore, gRNAs targeting the promoter region, specifically Spg1, Spg2, and Spg3, showed higher editing efficiencies (**Table 4**; **Figure 3B, C**), potentially due to factors such as chromatin accessibility, gRNA activity, or gRNA expression levels. Overall, ZmSpRYi in sorghum achieved editing efficiencies comparable to those reported in other crops, such as rice [19,20]. Given its ability to introduce larger deletions and to target AT-rich regions with greater flexibility, ZmSpRYi serves as a highly effective tool for promoter editing.

**Table 4.** Cis-regulatory element and gene exon editing efficiency of SpRY on phytoene desaturase and cytochrome P450 genes.

| Target | Rep.<sup>†</sup> | #Reg.<sup>‡</sup> | #Edits<br>%Efficiency<sup>§</sup> |  | %gRNA1<br>efficiency<sup>§</sup> |  | %RNA2<br>efficiency<sup>§</sup> |  | %gRNA3<br>efficiency<sup>§</sup> |  | %gRNA4<br>efficiency<sup>§</sup> |  | %gRNA5<br>efficiency<sup>§</sup> |  | %gRNA6<br>efficiency<sup>§</sup> |  |
| --- | --- | --- | --- | --- | --- | --- | --- | --- | --- | --- | --- | --- | --- | --- | --- | --- |
| *SbPDS* | 1<sup>st</sup> | 14 | 8 | 57.1 | 8 | 57.1 | 8 | 57.1 | 6 | 42.9 | 3 | 21.4 | 3 | 21.5 | 5 | 35.7 |
|  | 2<sup>nd</sup> | 50 | 23 | 46.0 | 21 | 42.0 | 22 | 44.0 | 18 | 36.0 | 11 | 22.0 | 14 | 28 | 16 | 32.0 |
| *SbCYP79A1* | 1<sup>st</sup> | 40 | 19 | 47.5 | 9 | 22.5 | 12 | 30.0 | 19 | 47.5 | 0 | 0 | NA<sup>¶</sup> | NA<sup>¶</sup> | NA<sup>¶</sup> | NA<sup>¶</sup> |

<sup>†</sup>Rep.= Replication
<sup>‡</sup>#Reg.= Regenerant number
<sup>§</sup>%gRNA efficiency = (# Edited plants divided by # plants + for hph) x 100
<sup>¶</sup>NA = not applicable

### *Cis*-regulatory element editing via ZmSpRYi in *CYP79A1* promoter

To test the potential of ZmSpRYi for promoter editing, we focused on *CYP79A1*, a key gene in the cyanogenic dhurrin biosynthesis pathway [51,52]. *CYP79A1* encodes a tyrosine N-monooxygenase that catalyzes the dedicated step in dhurrin production [53]. While complete knockout of this gene eliminates dhurrin toxicity, it also removes dhurrin’s benefit as a nitrogen reservoir and natural defense compound [54]. Modulation of *CYP79A1* expression via promoter editing would offer an approach for optimizing the agronomic potential of sorghum for diverse applications.

To explore the generation of diverse promoter edits, we examined sequence conservation of CYP79A1 promoter orthologs across C4 grasses, including *Sorghum bicolor* subspecies *drummondii* and *verticilliflorum*, as well as *Sorghum propinquum*, *Saccharum officinarum × spontaneum*, *Miscanthus sinensis*, and *Andropogon gerardii* (**Figure 3D**). Based on variation within the core promoter region, 1 kb upstream of the translational start site, compared to sorghum genotype RTx430, we designed four gRNAs targeting the *CYP79A1* promoter and 5′ untranslated region (UTR) at positions 600 bp, 383 bp, 212 bp, and 23 bp upstream of the translational start site (**Figure 3D**). These sites were selected with the aim of disrupting putative cis-regulatory elements (**Figure 3E**). ZmSpRYi and all four gRNAs, Cpg 1, 2, 3, and 4, were transformed into sorghum. Out of 40 independent T transformants, 19 were successfully edited at least by one gRNA, resulting in an overall editing efficiency of 47.5% (**Table 4**). Interestingly, the editing patterns varied among the four gRNAs. Of the four gRNAs, Cpg2 and 3 demonstrated the highest editing efficiencies, at 30.0% and 47.5%, respectively. In contrast, Cpg1 had 22.5% editing, while Cpg4 gave rise to 0% editing (**Table 4**). Our current findings demonstrate the feasibility of using ZmSpRYi-mediated promoter editing to attempt developing varieties with varying dhurrin levels.

## Discussion

Development of efficient, targeted, and versatile genome-editing tools is needed to advance genetic engineering and genome editing in bioenergy crops like sorghum. As a C4 plant, sorghum is naturally efficient in photosynthesis but additionally is highly resilient to environmental stresses, such as drought, flooding, and salinity, which could increase with climate changes [3,55]. However, the molecular mechanisms underlying such key agronomic traits and the ability to validate appropriate genetic resources to confirm functions remain significant challenges. To address these challenges, while introducing desirable genetic variations in sorghum, we developed a shuttle-binary vector toolbox to address current limitations in genome editing. This system simplifies genetic engineering by incorporating into the transformation construct the morphogenic gene *Wus2*, which accelerates more effectual delivery of editing machinery [11]. Additionally, several optimizations were also introduced to improve efficiency of CRISPR/Cas9-based genome editing in sorghum, including a codon-optimized Cas9 containing an intron, high-efficiency endogenous U3/6 promoters, and monocistronic gRNA expression cassettes. These modifications addressed previous limitations and significantly expanded the potential applications of this genome editing technology in sorghum.

### Wuschel integration simplifies the binary vector and improves transformation frequency

Previous studies demonstrated that the morphogenic regulators, BBM and WUS2, significantly enhance transformation frequency in monocot species, including sorghum [8,9]. However, continued expression of the morphogenic regulators often results in developmental aberrations during the vegetative stage. To address this issue, the CRE/LoxP recombination system was employed to excise the morphogenic genes, following somatic embryo initiation [9]. While effective, this approach requires increasing the binary vector size, complicating molecular cloning and construct development. Several approaches were used to overcome this challenge [8], including reducing construct size and development of systems that combine transient and stable *Wus2* expression [11,56]. In our approach, we redesigned the binary vector into a versatile delivery platform by incorporating an LR recombination site, thereby ensuring compatibility with the Gateway system, which is widely used for construct development among the plant research community. When paired with a *Wus2* expression cassette, we used this system to achieve transformation frequencies from 18.2% to 31.4% (**Table1 and 2**), rates comparable to those attained using the BBM/WUS2-assisted transformation approach [8].

Despite this success, limitations persisted with integration-based delivery of *Wus2*. Approximately 10% of the T_0_ transformants exhibited developmental deficiencies, such as curly leaves during the vegetative stage, potentially caused by ectopic expression of WUS2. To address this issue, pGL216 can be used as a platform for testing alternative morphogenic regulator genes, including *Wuschel* orthologs from other species or members of the *Wuschel* homeobox gene family (**Figure 1C**). These alternative candidate genes may enhance somatic embryo initiation while minimizing developmental disruptions, offering a more refined approach to transformation and regeneration.

### Optimizing *Cas9* and guide RNA expression cassettes increases editing efficiency

In our previous work, we successfully achieved *PDS* knockouts in sorghum using CRISPR/*Cas9*, with an editing efficiency of 16.7% [8]. However, enhancing genome-editing efficiency further requires overcoming certain challenges, particularly ensuring robust expression of the editing machinery, including the *Cas9* protein and gRNAs [14]. To address the editing challenges, we used a constitutive promoter to drive expression of the intronized maize codon-optimized nuclease genes, including *Cas9* and *SpRY*. Intronization was shown to enhance protein abundance at both transcriptional and translational levels in Arabidopsis [22]. For gRNA expression optimization, we utilized the sorghum endogenous snoRNA U6 promoters, which lead to high gRNA expression levels [23].

Previous studies suggest that transcriptional configurations, including polycistronic and monocistronic approaches, may affect editing efficiency [38]. However, it remained unclear whether a polycistronic configuration compromises editing efficiency in sorghum. To investigate this, we conducted a side-by-side comparison of polycistronic and monocistronic gRNA transcriptional configurations using Cas9 and SpRY nucleases. Results indicate that the monocistronic gRNA configuration led to significantly higher editing efficiency, up to 97.6% with Cas9 and 88.0% with SpRY, compared to 61.7% and 26.6% respectively observed with the polycistronic configurations (**Tables 1 and 3**). Additionally, gRNA1.1 exhibited lower editing efficiency in the polycistronic configuration compared to the monocistronic approach, suggesting that the temporal processing of the first exogenous tRNA in the polycistronic construct may be suboptimal in sorghum (**Figure 2D**). We also observed, when using the polycistronic constructs, a delayed onset of albino phenotypes resulting from *PDS* knockouts. This delay further supports the notion that transcriptional processing inefficiencies might contribute to the reduced editing efficiency observed with polycistronic gRNA configuration in sorghum. To improve editing outcomes, we optimized the monocistronic configuration for gRNA expression using a collection of monocot-derived snoRNA promoters. This collection includes U6 promoters from sorghum, wheat, and maize, as well as U3 promoters from wheat and rice (**Figure S1**), all of which have been experimentally validated in sorghum (**Figure 3B, Figure S5**). Promoters were incorporated into Level 0 constructs, providing flexible tools for expressing multiple gRNAs independently. This modular approach facilitates multiplex genome editing and is particularly well-suited for applications such as promoter editing and other complex genetic modifications.

To evaluate the impact of temperature on editing efficiency, we conducted an editing experiment using different incubation temperatures, 25°C and 28°C, following Agrobacterium infection. Interestingly, no apparent difference in editing efficiency was observed between the two temperatures (**Table 1**). This outcome may be attributed to the high baseline editing efficiency achieved through the modifications introduce into both the Cas9 and gRNA components. Elevated temperatures did not confer further improvements, probably because the optimized CRISPR/Cas9 constructs were already near their maximal efficiency. In contrast, tissues incubated at 28°C exhibited significant necrosis; whereas those maintained at 25°C remained healthier and produced more regenerants on average (**Figure S3**). Therefore, temperatures above 28°C were excluded from further experimentation. Our findings suggest that 25°C is a more suitable incubation temperature for achieving efficient genome editing in sorghum.

### Expanding the scope of genome editing using ZmSpRYi

One potentially impactful application of genome editing in crops is creating cis-regulatory variations to diversify genetic resources for improving desirable traits [44]. Editing of cis-regulatory elements in promoters and UTRs, which are typically AT-rich, is often constrained when using Cas9, due to its strict NGG PAM requirement and in sorghum its tendency to produce small deletions. To address the need for AT sequences, we considered several potential genome editing candidates to target AT-rich regions, including Cas9 variants (Cas9-NG, xCas9, and SpRY) and other Cas orthologs (Cas12a and Cas12b), which exhibit reduced dependency on CG content at the PAM site [24,57–60]. Among these editing candidates, we chose SpRY which has the highest flexibility due to its PAM-less feature [24].

Using the same gRNAs that targeted the PDS gene in our previous CRISPR/*Cas9*-mediated editing experiments [8], in the present study Zm*SpRY*i achieved an editing efficiency of 88.0% (**Table 3**). However, previous studies suggested that use of *SpRY* results in compromised double-strand break activity, particularly with non-NGG PAM sequences [19,20]. To further investigate ZmSpRYi activity with non-NGG PAM sequences, we tested its performance in combination with WUS2. Previous research indicated that integration of *Wus2* can enhance CRISPR/*Cas9*-mediated gene dropout frequency [11]. However, it remained unclear as to whether *Wus2* integration can similarly enhance SpRY activity. To address this question, we designed six gRNAs with NAA, NAT, NTA, and NTT PAM sequences (**Figure 3A**). Across 64 independent transformants, an overall editing efficiency of 48.4% was observed (**Table 4**), which is comparable to the editing efficiency reported in rice [19,20]. Interestingly, regardless of the PAM sequences, gRNAs targeting promoter regions (48.4%, 46.9%, and 37.5%) exhibited higher editing efficiencies than those targeting coding regions (21.9%, 26.6%, 32.8%) (**Table 4**), probably due to differences in chromosome accessibility among these genomic regions.

Another concern for SpRY is the potential for off-target effects, including self-editing, which refers to mutagenesis occurring within the gRNA expression cassette [19,20], possibly leading to unintended edits. To assess this risk in edits with high knockout scores, we amplified and sequenced potential off-target sites, including regions within the T-DNA. We found that 6.7% (1/15) of plantlets from polycistronic configurations targeting PDS gRNAs with NGG PAMs showed self-editing, whereas monocistronic configurations exhibited self-editing frequencies of up to 33.3% (6/18) within the T-DNA (**Figure S6**). Moreover, gRNAs with non-canonical PAMs showed 66.6% self-editing frequency on at least one gRNA, probably due to the increased number of gRNAs, up to six in the construct (**Figure S6**). The target regions were also successfully edited in these cases, suggesting that self-editing occurred either concurrently or sequentially with the targeted editing process. Taken together the present study highlights the promise of *SpRY* for expanding the genome editing toolbox in AT-rich regions, while acknowledging challenges like compromised activity and self-editing risks.

### Potential of ZmSpRYi for efficient promoter editing

Results of the present study demonstrates the potential use of SpRY, a PAM-flexible Cas9 variant, as a tool for promoter editing in plants. For example, by editing the CYP79A1 promoter, it should be possible to generate quantitative variations to limit dhurrin production to increase the safety of sorghum as livestock feed. Toward this end, the SpRY construct achieved a moderate overall editing efficiency of 47.5%, with over half of the edited plants showing a knockout score exceeding 50% at the second and third gRNA sites (**Table 3**). These editing results suggest SpRY is a viable tool for editing AT-rich regulatory regions, where conventional Cas9 is restricted by stringent PAM requirements. Furthermore, combining adjacent gRNAs, such as Spg2 and Spg, enables the targeting of potential cis-regulatory elements and may promote larger dropouts, due to the PAM flexibility of SpRY.

Despite its promise, SpRY remains under-optimized. The moderate editing rate suggests that SpRY’s PAM flexibility may compromise target engagement, possibly due to reduced DNA binding affinity or inefficient R-loop formation [49]. Artificial intelligence-assisted optimization represents a strategy to enhance the performance of PAM-less Cas9 variants, like SpRY, and further alleviate PAM-related constraints [61]. While SpRY expands the range of targetable sequences, gRNA performance was inconsistent. Although *in silico* predictions using certain algorithms indicated that all four gRNAs should be highly efficient, only the second and third showed moderate activity, while the fourth was non-functional. This discrepancy underscores the limitations of current gRNA efficiency prediction tools, which fail to consider biological variables, such as chromatin accessibility especially near transcription start sites [62]. Refining gRNA design algorithms that incorporate epigenomic features and consider the activity of engineered editors, for example SpRY, might improve future targeting accuracy and efficiency [63].

## Concluding remarks

In the current study, we described high-efficiency transformation and editing approaches incorporating *wus2* integration with a robust Cas9-based genome editing toolkit, featuring intronized ZmCas9 and ZmSpRY. This platform is a versatile means for delivering genome editing machinery that can include base editors and prime editors. By integrating *wus2* with the Cas9 variant, SpRY, we expanded the editing scope to increase other desirable applications, such as promoter editing. These advancements provide a streamlined approach to generating diverse genetic variation in sorghum, facilitating crop improvement efforts to address food security challenges and contribute to carbon capture initiatives that address climate change.

## Materials and Methods

### Plant material and growth conditions

Seeds of *Sorghum bicolor* genotype, RTx430, were sourced from the Germplasm Resources Information Network (GRIN, USDA-ARS, PI 655998). Plants were cultivated in three-gallon pots containing SuperSoil (Rod Mclellan Company) in UC Berkeley’s Oxford Research Unit South Greenhouse, under controlled conditions of 30°C/24°C (day/night) with a 16-hour light/8-hour dark photoperiod. Immature embryos (IEs) were harvested 12–14d post-anthesis.

### Construct creation

To create pGL206 (**Figure 1C**), the *hygromycin B phosphotransferase* gene (*hph*), driven by the *Panicum virgatum Ubiquitin2* promoter (PvUbi2p), was amplified from pANIC10A [64] using the primer SorEd1 and 2 (**Table S1**). All primers used for construct building are described in **Table S1.** The terminator for sorghum PEPC was amplified from pPHP85425 [8] using the primers SorEd3 and 4. These two parts were assembled by overlapping PCR. The vector backbone of pPHP85425 was amplified using the primers SorEd5 and 6. In addition, the Buffer3 and attR4-attR3 fragments were amplified from pPHP85425 using the primers SorEd7 and 8. These three fragments were assembled using Gibson assembly (NEBuilder® HiFi DNA Assembly Master Mix). To make the construct pGL216 (**Figure 1C**), the *ZsG* expression cassette was amplified from pGL190 [8] using the primer SorEd9 and 10. The amplified cassette was then inserted into pGL206 through *Sfi*I digestion and subsequent ligation. To create pGL222 (**Figure 1C**), the partial backbone of pPHP96564 [11] and the *Wus2* expression cassette were amplified from pPHP96564 using the primers SorEd11 and 12. The attR4-attR3 region was amplified from pPHP85425 using the primers SorEd13 and 14. The *hygromycin B phosphotransferase* (*hph*) gene was amplified from pGL206 using the primers SorEd15 and 16. The rest of the backbone of pPHP96564 was amplified with the primers SorEd17 and 18. These fragments were reassembled into the final vector using Gibson assembly. Additionally, the *ZsG* gene, driven by the *Sorghum bicolor ubiquitin1* promoter, was amplified from pGL190 with the primers SorEd19 and 20. The ZsG expression cassette was digested with *Nhe*I and *Pme*I and ligated into the construct to finalize the construct, pGL222, validated using Nanopore sequencing.

To optimize shuttle constructs, the sorghum U6.2.3 (SbU6.2.3) promoter [23] from sorghum genotype RTx430 was amplified using the primers SorEd21 and 22. pGL198 [8] was linearized with primers, SorEd23 and 24. The OsU3 promoter was replaced by SbU6.2.3 using Gibson assembly. Additionally, ZmCas9 was replaced by ZmCas9i using primers SorEd25 and 26 for vector linearization and primers SorEd27 and 28 for ZmCas9i amplification from pPHP96564, resulting in the shuttle vector, pGL220. Based on the polycistronic shuttle vector construction, the monocistronic shuttle vector was created by inserting two gRNA expression cassettes by Gibson assembly with primers SorEd29-36 to create the monocistronic expression cassettes. Cas9 and gRNA polycistronic and monocistronic cassettes were transferred to pGL222 to create binary constructs pGL220 and 221 by LR recombination. To create pGL224, we used the same strategy as described above to replace the OsU3 promoter and ZmCas9 with the SbU6.2.3 promoter and ZmCas9i. To streamline construct creation, level 0 constructs were developed. These constructs were created by fusing the gRNA scaffold with various snoRNA promoters (including sorghum U6_2.3 [SbU6_2.3], sorghum U6_3.1 [SbU6_3.1] [23], rice U3 [OsU3] [65], maize U6 [ZmU6] [66], wheat U3 [TaU3] [67], and wheat U6 [TaU6]) [68] using overlapping PCR with primers SorEd49 through 62. Fused DNA fragments were cloned into a pJET1.2 vector using CloneJET PCR cloning kit (ThermoFisher Scientific) and validated by Sanger sequencing. These level 0 constructs served as templates for creating both polycistronic and monocistronic gRNA editing constructs (**Figure S2**).

To create the ZmSpRYi construct, pGL224 was used as the template, and a total of 14 nucleotide mutations were introduced. The first set of mutations on the first exon of ZmCas9i targeted the removal of a BsaI recognition site and the substitution of alanine 61 with arginine. These nucleotide mutations were incorporated using primers SorEd37 and SorEd38, enabling the amplification of two overlapping fragments with primers SorEd39 and SorEd40. The overlapping fragments were assembled through overlapping PCR. The modified sequence replaced the original sequence via BamHI and BspEI digestion, followed by ligation. The remaining ten mutations were introduced using a synthetic oligonucleotide chemically synthesized by Genewiz (http://genewiz.com). All these mutations were inserted into the 3′ end of ZmCas9i through Gibson assembly. Primers SorEd41 and SorEd42 were used to amplify the insert, while primers SorEd43 and SorEd44 were employed for vector linearization. Additionally, two unwanted BsaI restriction sites were eliminated via primer-directed mutagenesis using primers SorEd45 through SorEd48, resulting in the construct, pGL225 (**Figure S1**). Designed for Agrobacterium-mediated transformation, the ZmCas9i gene in pGL220 and pGL221 was replaced with the ZmSpRYi gene to create pGL226 and pGL227. To generate the pGL236 construct targeting the *PDS* promoter and exons, gRNAs were designed using the CRISPOR tool [69]. For gRNAs ending with NAN and NTN motifs, we amplified both the gRNA scaffolds and the snoRNA promoter from level 0 constructs using primers that incorporated the guide sequences. These primers also served as adaptors for overlapping PCR. The primers used for assembling the six gRNA expression cassettes are listed as SorEd63-72.

The assembled PCR product was digested with *Bsa*I and ligated into the *Bsa*I-digested vector pGL225. The integrity of the construct was validated via Nanopore sequencing and subsequently recombined into the binary vector pGL222 through LR recombination, resulting in the final construct, pGL236, to be used for Agrobacterium-mediated transformation.

For the pGL237 construct targeting the *CYP79A1* promoter, gRNAs were designed based on conserved promoter and untranslated regions identified across homologs from the sorghum genus, *Miscanthus sinensis*, *Saccharum officinarum*, and *Andropogon gerardi* (**Figure 3D**). The CRISPOR tool was used to design gRNAs with the goal of disrupting transcription by targeting predicted cis-regulatory elements. Considering the distribution of cis-elements and optimal gRNA spacing, we selected gRNAs spaced approximately 200 bp apart within the promoter region (**Figure 3D**). The gRNA expression cassettes were amplified using primers SorEd73-78, assembled via overlapping PCR and ligated into pGL225, as previously described. The construct was verified with Nanopore sequencing and finalized by LR recombination to generate pGL237 for further Agrobacterium-mediated transformation.

### Agrobacterium-mediated transformation

The binary constructs were introduced into an auxotrophic strain of *Agrobacterium tumefaciens* LBA4404 Thy- <u>(Anand et al. 2017)</u> by electroporation. Transformed Agrobacterium strains were plated on yeast extract peptone (YEP) medium supplemented with 100 mg/L thymidine, 50 mg/L gentamicin, and 50 mg/L spectinomycin. Details on the storage and preparation of the Agrobacterium strains followed a previously published protocol [70]. In summary, Agrobacterium stocks were streaked onto YEP plates containing appropriate antibiotics to generate master plates. After a 2-day incubation at 28°C, 5 to 6 colonies from the master plates were selected to establish working cultures on YEP plates. Following overnight incubation at 28°C and just prior to inoculation of sorghum immature embryos, Agrobacterium suspensions were prepared in PHI-I medium [71], with optical density (OD550 nm) adjusted to 0.7; silwet L-77 (0.005% v/v) and acetosyringone (0.2 mM) were added to finalize the Agrobacterium inoculation medium. IEs were inoculated with Agrobacterium suspension culture for 5 minutes, then placed on co-cultivation medium at 25 °C for one week. After co-cultivation, the IEs were transferred to resting medium for an additional week and then moved to either 25 °C or 28 °C. Selection followed on embryo maturation medium containing 15 mg/L hygromycin until regeneration commenced. Regenerated plantlets at 1–2 cm in height were then transferred to rooting medium until fully rooted (**Figure 1C**). Transformation frequency was calculated as the percentage of IEs that successfully produced transformants.

### Visualization of T-DNA integration

The ZsGreen reporter gene was used as a visible marker to track T-DNA integration and segregation. To monitor T-DNA integration following Agrobacterium-mediated inoculation, transformed IEs were examined weekly using a Lumar v12 epifluorescence stereo microscope and a QImaging Retiga-SRV camera (Zeiss, UC Berkeley Biological Imaging Facility). Response of IEs to WUS expression was assessed by analyzing the number and size of fluorescent tissues. Prior to transferring shoots to rooting medium, regenerants exhibiting fluorescence were selected for further genotyping and phenotyping.

### Genotyping of editing events

To test putative T_0_ transgenic lines for successful editing of target genes, genomic DNA from young sorghum leaves was isolated using a urea buffer as described [8]. For the target regions in putative PDS edits due to gRNA1.1 and 2.2, primers SorEd79 and 80 were employed. To genotype editing by non-canonical gRNAs on the PDS gene, primers SorEd81 and 82 were used. For genotyping CYP79a1 cis-regulatory elements, primers SorEd83 and 84 were used for PCR amplification. Amplicons were amplified and then purified with the GeneJet PCR Purification Kit (Thermo Scientific). Purified PCR products were Sanger-sequenced at the UC Berkeley DNA Sequencing Facility, using the primers for PCR amplification. Sequencing data were analyzed with the Inference of <u>C</u>RISPR <u>E</u>dits (ICE) tool [43], and knockout scores, referring to proportion of indels that indicate a frameshift or are more than 21 bp in length, were used to assess editing efficiency and the likelihood of transmission to the next generation. Editing efficiency represents the percentage of plant cells within harvested somatic samples that successfully undergo knockout events, as determined by the knockout score.

### Phenotypic characterization of T_0_ PDS plants

T_0_ plantlets were phenotyped while growing in rooting media. Each plantlet was scored for the percentage of albino phenotype, with 100% indicating plants that were completely albino and 0% indicating plants that were completely green. The phenotype score was compared with the knockout percentage obtained using the ICE tool to determine correlation between the two observations.

## Supporting information

Supplementary materials will be used for the link to the files on the preprint site

## Resource availability

### Lead contact

Requests for further information and resources should be directed to the lead contact, Peggy G. Lemaux

### Material availability

pGL constructs can be used for research purposes by signing Material Transfer Agreements with Corteva Agriscience and UC Berkeley. Using vectors for commercial applications requires a paid non-exclusive license agreement with Corteva Agriscience. The lead PI will share sequences and plasmids upon reasonable request.

### Data availability

All data generated in this work are reported in this manuscript. Any additional information will be made available from the lead contact upon reasonable request.

## Acknowledgments

This project was supported by the Chan Zuckerberg Initiative through the Innovative Genomics Institute and by the U.S. Department of Energy, Office of Science Energy Earthshot Initiative, as part of the Terraforming Soils EERC led by Lawrence Livermore National Laboratory under Award No. SCW1841. PGL acknowledges support from the US Cooperative Extension Service through the Division of Agriculture and Natural Resources of the University of California. DFS is an investigator in the Howard Hughes Medical Institute. JS acknowledges support from USDA-ARS Crop Improvement and Genetics Unit Albany, CA and ORISE Research Participation Program. We thank Dr. William Gordon-Kamm at Corteva Agriscience for helpful conversations and for the constructs, pPHP96564 and pPHP85425. We thank Dr. Roger Thilmony at USDA-ARS Crop Improvement and Genetics Unit Albany for informative discussions. We thank talented undergraduate students, Malia Ariyoshi, Amada Niemela, Emma To, Nessa Kmetec, Isabella Lombardo, Dasha Wohlfarth, and Sanay Parikh, for their help in this work. We are grateful for the help of the staff of the UC Berkeley Biological Imaging Facility and the UC Berkeley Oxford Tract Greenhouse Facility.

## Author contributions

Conceptualization: J.S., K.A., E.G., D.S. and P.G.L.. Funding acquisition: D.S. and P.G.L.. Methodology: J.S., K.A., and S.A.. Investigation: J.S., K.A., S.A., and M.R.. Formal analysis: J.S. and K.A.. Visualization: J.S., M.R., and K.A. Writing—original draft: J.S., K.A., and T.M.. Writing—review and editing: J.S., K.A., S.A., T.M., and P.G.L.. Supervision: J.S., D.S. and P.G.L.

## Declaration of interest

Manuscript authors have no relationship with Corteva Agriscience. The authors declare that they have no competing interests.

## Notes

### Competing Interest Statement

The authors have declared no competing interest.

### Summary of Updates

Author list and affiliations updated. Figure 1 and 2 in the version 1 merged to Figure 1. Figure 3 and 4 in the version 1 merged into Figure 2. Figure 3 a-c revised to show the PDS editing results using SpRY and gRNAs with non-canonical PAMs. Figure 3 d-f added to demonstrate SpRY application on the promoter editing of dhurrin biosynthesis gene, CYP79A1. Table 1-4 revised to update the latest results. Supplementary figure 4-5 updated to show the mutations to create SpRY, deletion position and indel size, as well as self editing rate. Section on SpRY editing using gRNAs with non-canonical PAMs updated to clarify the editing efficiency of SpRY. Section on CYP79A1 promoter editing updated to add a case of SpRY editing. Section on self editing updated to add more detailed data. The Acknowledgements section has been updated to more accurately reflect detailed funding sources.

