## Supplementary materials will be used for the link to the files on the preprint site for "High-frequency sorghum transformation toolkit enhances *Cas9* efficiency and expands promoter-editing capability with *SpRY*"

**Supplementary figures**

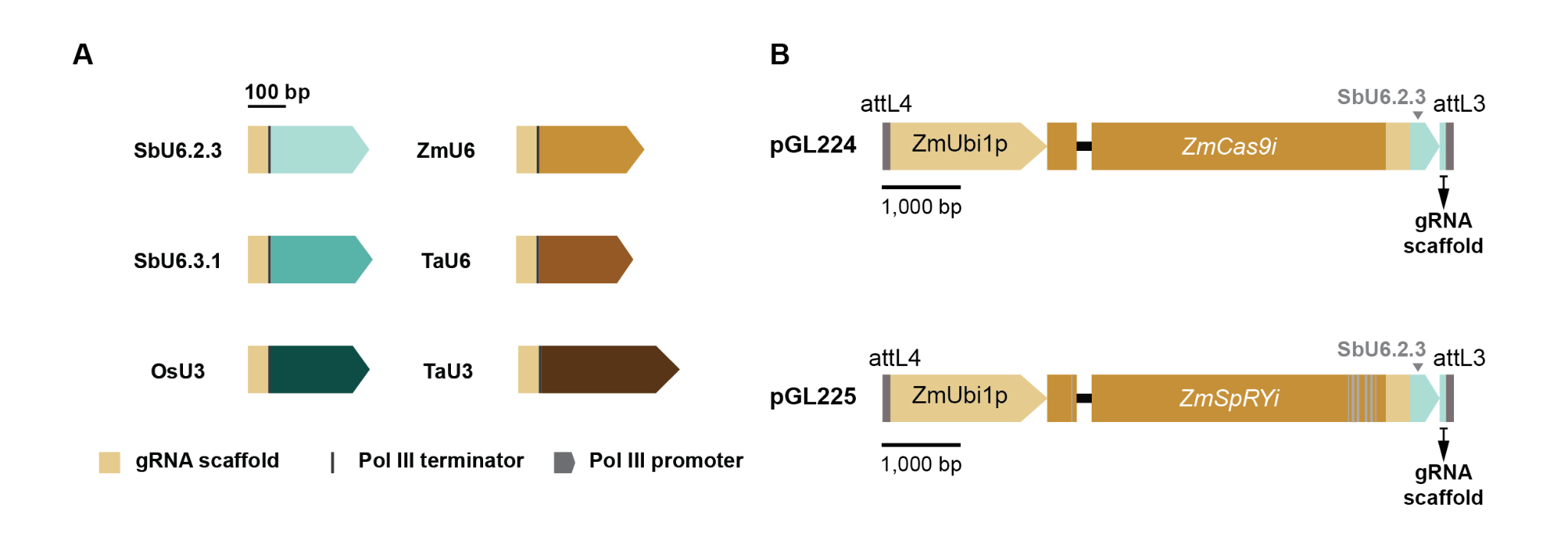

**Figure S1. Maps of level 0 constructs and shuttle vector backbones. (A)** Maps of level 0 constructs. Gray rectangles indicate gRNA scaffold. Rectangles with arrows indicate different Pol III promoters. Dark gray vertical lines indicate terminators of gRNA expression cassettes. **(B)** Maps of shuttle vector backbones. Rectangles with arrows indicate promoters. Other rectangles indicate exons (dark-colored) or terminators (light-colored). Black horizontal lines indicate introns. Grey vertical lines indicate Gateway recombination sites.

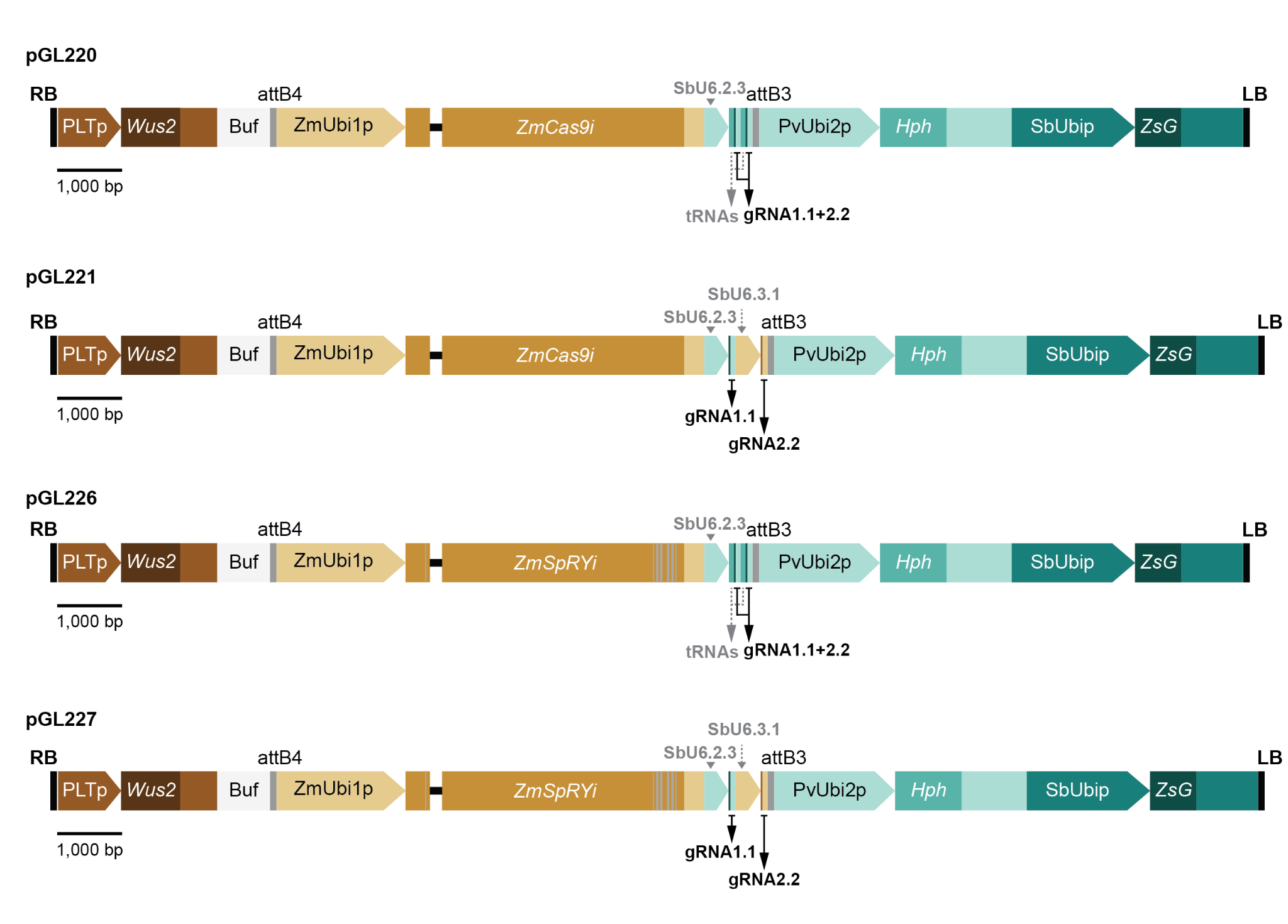
**Figure S2. Maps of the binary constructs used for Agrobacterium-mediated transformation.**

Vector maps for CRISPR-mediated genome editing in sorghum. **pGL220**: Binary vector for CRISPR/Cas9 editing, incorporating polycistronic gRNA expression cassettes driven by the SbU6.2.3 promoter. **pGL221**: Binary vector for CRISPR/Cas9 editing, utilizing monocistronic gRNA expression cassettes driven by the SbU6.2.3 and SbU6.3.1 promoters. **pGL226**: Binary vector for CRISPR/SpRY editing, featuring polycistronic gRNA expression cassettes driven by the SbU6.2.3 promoter. **pGL227**: Binary vector for CRISPR/SpRY editing, employing monocistronic gRNA expression cassettes driven by the SbU6.2.3 and SbU6.3.1 promoters. All gRNAs target the sorghum PDS gene. Rectangles with arrows represent promoters. Solid rectangles indicate exons (dark-colored) or terminators (light-colored); light gray rectangles denote buffer regions. Black vertical lines mark T-DNA borders, and gray vertical lines indicate Gateway recombination sites.

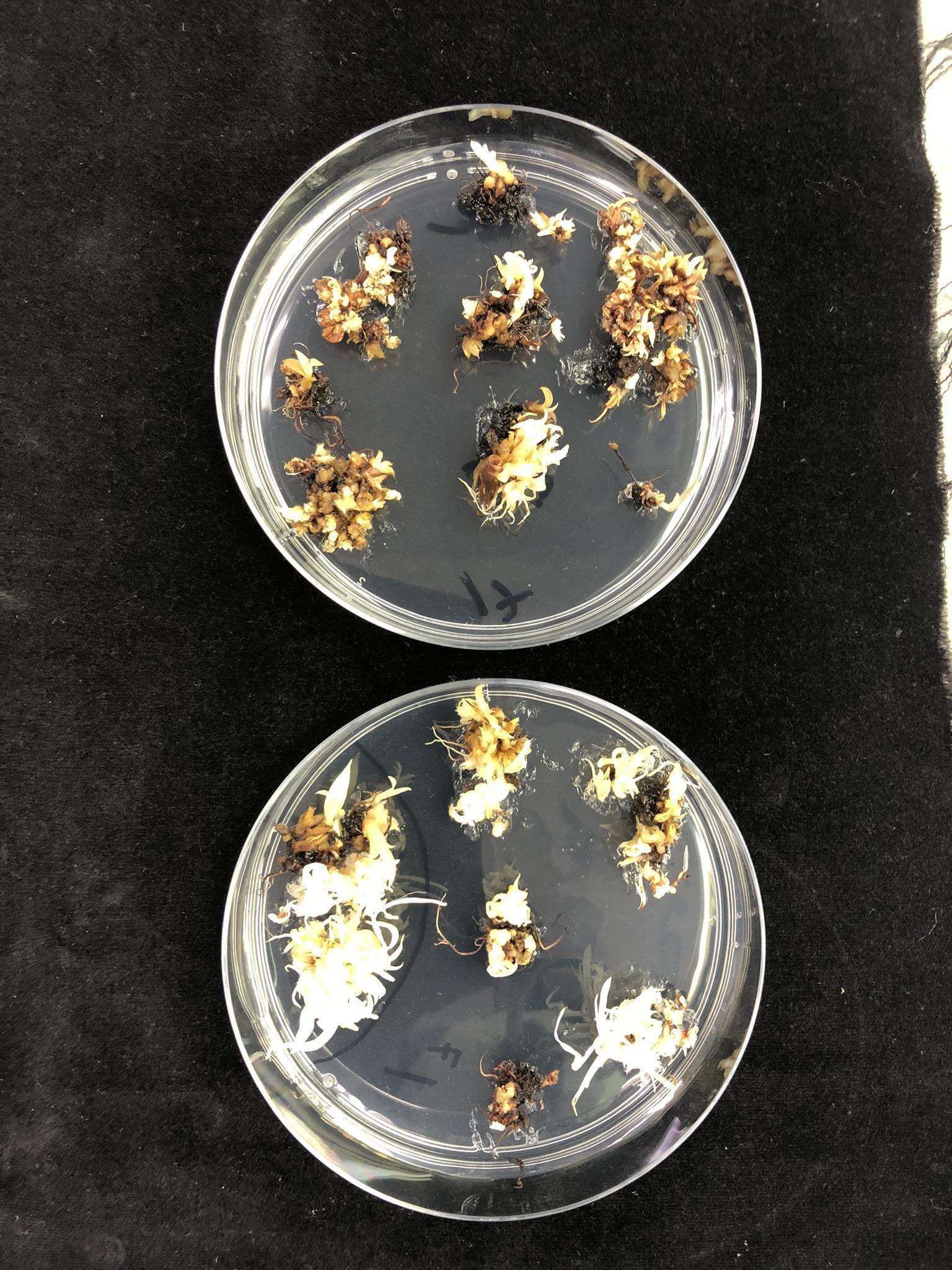

**Figure S3**. pGL220 transformed tissues on EMM. Tissues kept at 28^o^C (left) showed more necrosis (brown tissue) compared to those kept at 25^o^C (right).

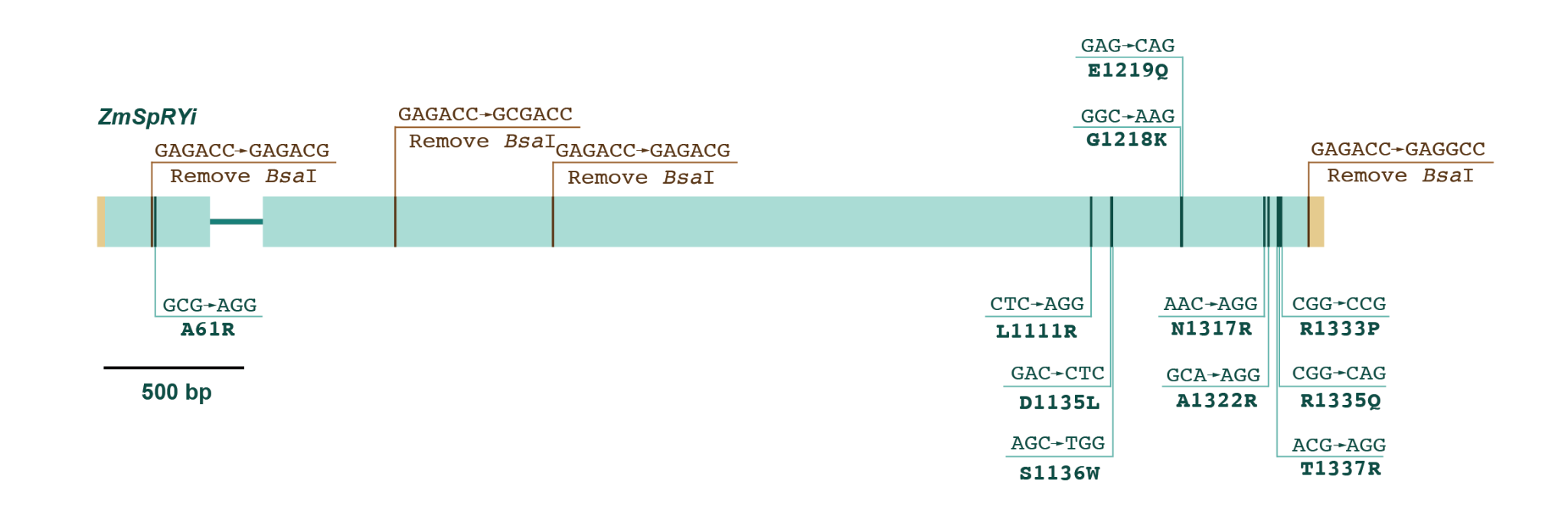

**Figure S4.** Mutagenesis of ZmCas9 to generate ZmSpRYi. *Bsa*I restriction sites were removed through site-directed mutagenesis (indicated by red vertical lines). Eleven amino acids were altered by primer-directed mutagenesis (indicated by green vertical lines).

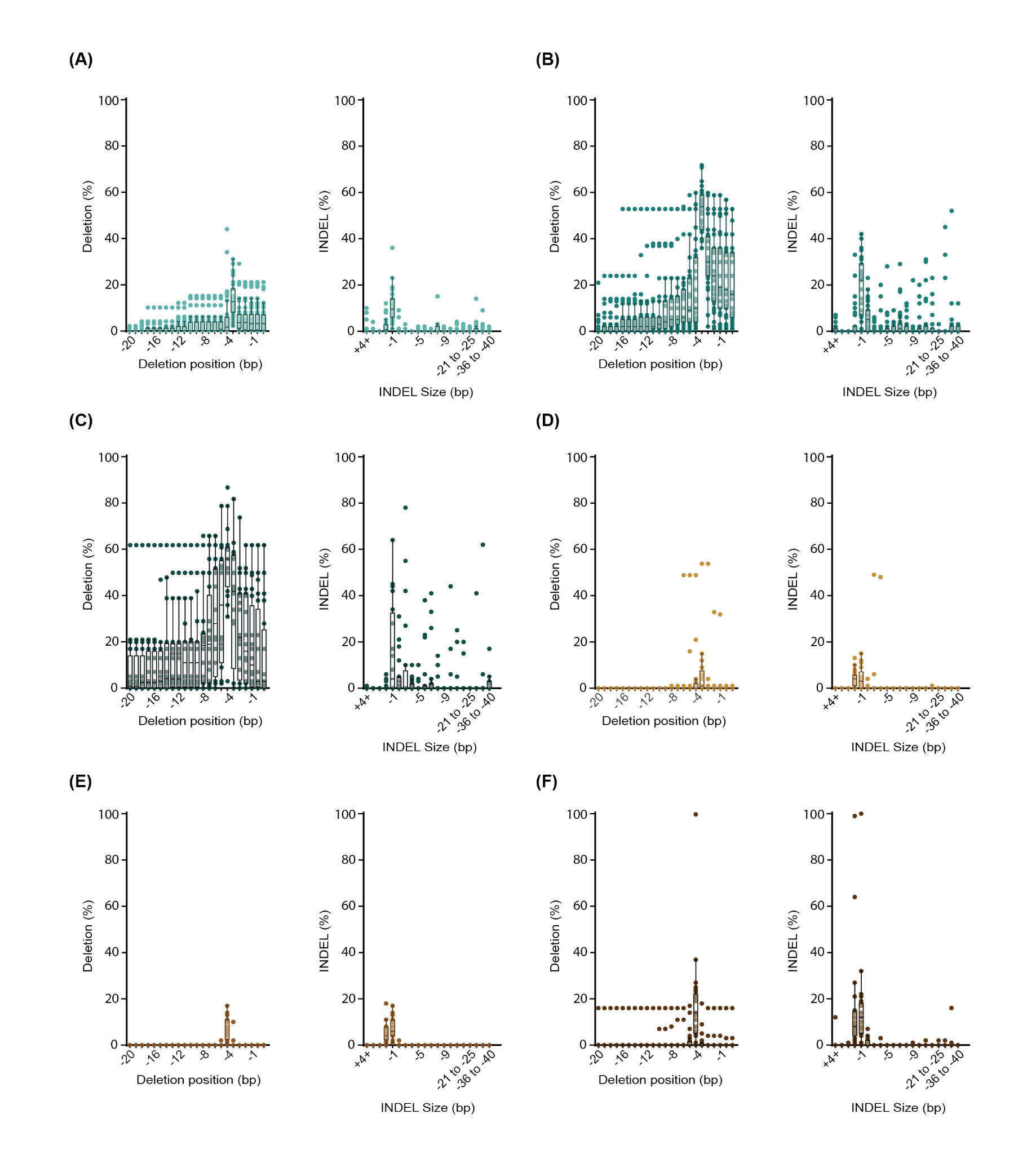

**Figure S5**. Deletion position and indel size of the non-canonical gRNAs. SbPDS gRNAs (Spg) targeting the promoter, 5’UTR and exons of SbPDS gene. Spg1 (A), Spg2 (B), Spg3 (C), Spg4 (D), Spg5 (E), and Spg6(F). Each dot represents a biological replicate. The distribution of deletion and indel percentage from non-canonical gRNAs are shown by highlighting the median and quartiles of each gRNA. For ZmSpRYi-mediated editing with non-canonical gRNAs, n=31.

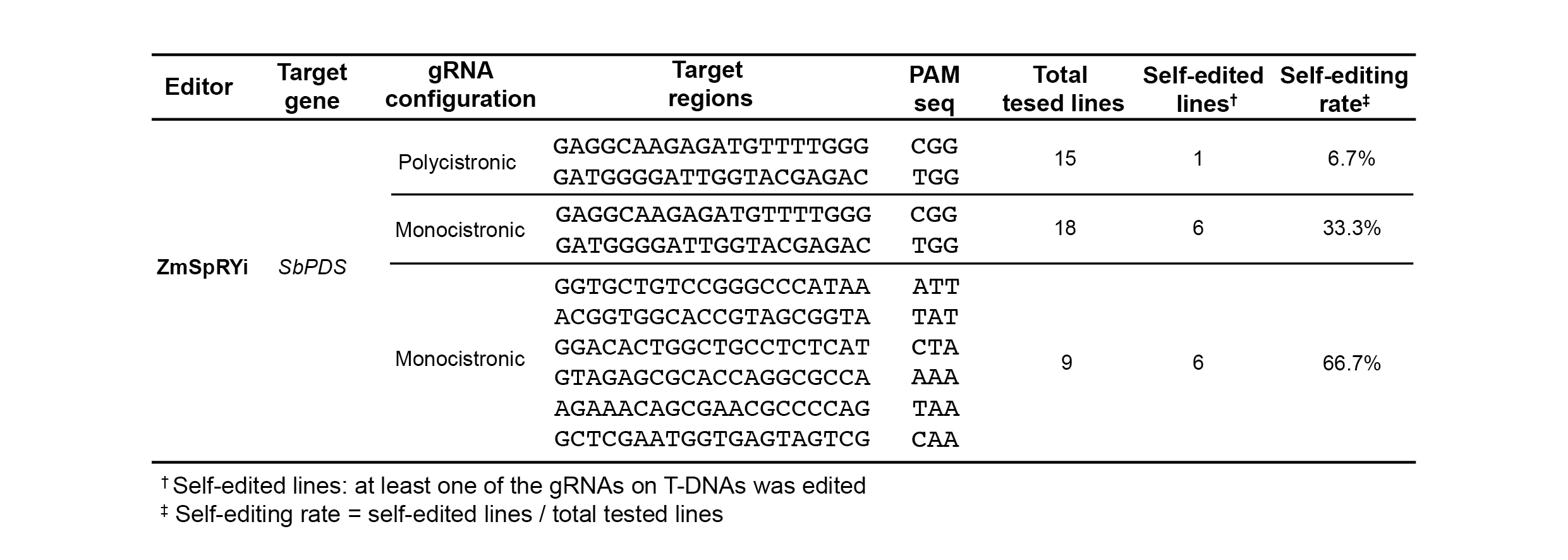

**Figure S6.** Self-editing rates observed in the T_0_ generation of SbPDS edited lines using ZmSpRYi. Self-editing events were identified through PCR amplification, followed by Sanger sequencing and ICE analysis. A construct was considered self-edited when at least one gRNA target site within the T-DNA region showed evidence of editing (knockout score is over 5%).

**Supplementary tables**

**Table S1.** Primer list

| **Primer** | **Sequences 5'- 3'** |
| --- | --- |
| SorEd-1 | TCGAGTTTTTCAGCAAGATGAAGCCAACTAAACAAGACCATAACCATGGTGACAT |
| SorEd-2 | AGGATCTGGCCTATTTCTTTGCCCTCGGACGAGTGCT |
| SorEd-3 | AAGAAATAGGCCAGATCCTCGGTGTACAAATAACCCG |
| SorEd-4 | TGTAGGAGATCTTCTAGAAAGATTATGTTGCTTAGGCCTCTTTTTGCCACGG |
| SorEd-5 | AAGCTTCGGTCCGGGCCTAGAAG |
| SorEd-6 | GGTAAGTCCGGAACTATAGCTAAGTCGTACGCTACT |
| SorEd-7 | CTATAGTTCCGGACTTACCAGTCCTACTAGTTAGTTAGGCGATCGCCCT |
| SorEd-8 | TTGGCTTCAGATCCCAGGGTACCGAGCTCGAATTC |
| SorEd-9 | AGGCCTAGAAGGCCCTGCCCTTAAGGCCAATTGTTCAAGATTCATTC |
| SorEd-10 | AGGCCTAGAAGGCCGAAGCCTACCAAAGCAAAGCGTTTGTCG |
| SorEd-11 | GATCGAGCTGGCAGACAAAGCAATAACCCACACAG |
| SorEd-12 | GCGATCGCCGGATGATTAGTCGCCACAAAATGCTTATACTGATGT |
| SorEd-13 | GTGGCGACTAATCATCCGGCGATCGCGCTGACAAGCG |
| SorEd-14 | GGTCTTGTTTAGTTGGCTTCAGATCCCAGGGTACCGAGCTCGAA |
| SorEd-15 | CCCTGGGATCTGAAGCCAACTAAACAAGACCATAACCATGGTGACAT |
| SorEd-16 | TTCGGGCCGCATTTATGTTGCTTAGGCCTCTTTTTGCCACGGAATTTGAG |
| SorEd-17 | CTAAGCAACATAAATGCGGCCCGAATAGGGGGA |
| SorEd-18 | TTATTGCTTTGTCTGCCAGCTCGATCCAAGAGTCAACGTTTGTGCC |
| SorEd-19 | CGAGCTAGCCTGCCCTTAAGGCCAATTGTTCAAGATTCATTC |
| SorEd-20 | GTGTTTAAACGAAGCCTACCAAAGCAAAGCGTTTGTCG |
| SorEd-21 | CATGATTACGCCAAGCTTCAGTTTTTCCTACGTTCAATGCTCGTGCCGT |
| SorEd-22 | GAGACCTCGGTCTCCGAAGCAGCTGGTGCGCTCGCGA |
| SorEd-23 | GGAGACCGAGGTCTCGGTTTTAGAGCTAGAAATAGCAA |
| SorEd-24 | AAGCTTGGCGTAATCATGTCATAGCTGTTTCCTGT |
| SorEd-25 | TGTTTTGCGGCGCGCCCAGTTTTTCCTACGTTCAATG |
| SorEd-26 | CTAGAGCGGCCGCGGCCATACAAG |
| SorEd-27 | CTTGTATGGCCGCGGCCGCTCTAGATGGCACCGAAGAAGAAGCGCAAGGT |
| SorEd-28 | AAAAACTGGGCGCGCCGCAAAACACACCTAGACTAGATTTGTTTTGCTAACCCAATTGA |
| SorEd-29 | ATGGCGCGCCCAGTTTTTCCTACGTTCAATGCTCGTGCCGT |
| SorEd-30 | CCCAAAACATCTCTTGCCTCGAAGCAGCTGGTGCGCTCGCGA |
| SorEd-31 | CCAGCTGCTTCGAGGCAAGAGATGTTTTGGGGTTTTAGAGCTAGAAATAGCAAGTTAAA |
| SorEd-32 | GTTTTACAAGGGATGGCCGGAGCTCAAAAAAAAGCACCGACTCGGTGCCACT |
| SorEd-33 | TGAGCTCCGGCCATCCCTTGTAAAACTTTAATTTTTTTACTAGTA |
| SorEd-34 | GTCTCGTACCAATCCCCATCGATGCGGTGCCTGCGCCT |
| SorEd-35 | CCGCATCGATGGGGATTGGTACGAGACGTTTTAGAGCTAGAAATAGCAAGTTAAAA |
| SorEd-36 | AGTCCTAGGTTAATTAAAAAAAAAGCACCGACTCGGTGCCACT |
| SorEd-37 | AGCGGCGAGACGGCGGAGAGGACCAGGCTCAAGAG |
| SorEd-38 | CTCTTGAGCCTGGTCCTCTCCGCCGTCTCGCCGCT |
| SorEd-39 | CTAGGATCCATGGCACCGAAGAAGAAGCG |
| SorEd-40 | GTTTCTTCCGGAGGTGGTAGATTGTCGG |
| SorEd-41 | GGGCGGGTTCAGCAAGGAGTCCATCAGGCCGAAGCGCAACTCC |
| SorEd-42 | GTGGCCTCTTGTCTCCGCCCAGCTGG |
| SorEd-43 | GGGCGGAGACAAGAGGCCACGGGACCGCCA |
| SorEd-44 | GGACTCCTTGCTGAACCCGCCC |
| SorEd-45 | GGCCCAGATAGGCGACCAGTACGCG |
| SorEd-46 | GTTCCAGGGTGTGATCGTCTCCTCCGACTTC |
| SorEd-47 | GAAGTCGGAGGAGACGATCACACCCTGGAAC |
| SorEd-48 | CGCGTACTGGTCGCCTATCTGGGCC |
| SorEd-49 | GTTTTAGAGCTAGAAATAGCAAGTTAAAATAAG |
| SorEd-50 | AAAAAAAAGCACCGACTCGGTGCC |
| SorEd-51 | GCACCGAGTCGGTGCTTTTTTTTCAGTTTTTCCTACGTTCAATGCTCGTGC |
| SorEd-52 | GAAGCAGCTGGTGCGCTCG |
| SorEd-53 | GCACCGAGTCGGTGCTTTTTTTTGAGCTCCGGCCATCCCTTGTAAAAC |
| SorEd-54 | GATGCGGTGCCTGCGCCTC |
| SorEd-55 | GCACCGAGTCGGTGCTTTTTTTTAAGGAATCTTTAAACATACGAACAGATCACTTAAAG |
| SorEd-56 | AGCCACGGATCATCTGCACAAC |
| SorEd-57 | GCACCGAGTCGGTGCTTTTTTTTGCTGTTTTTGTTAGCCCCATCGA |
| SorEd-58 | AATTCGGTGCTTGCGGCTCG |
| SorEd-59 | GCACCGAGTCGGTGCTTTTTTTTGCATGAATCCAAACCACACGGAG |
| SorEd-60 | TCGTGCTTCTTGGTGCCGC |
| SorEd-61 | GCACCGAGTCGGTGCTTTTTTTTAAGCCCGTTATTCTGACAGTTCTGG |
| SorEd-62 | AAGTCTGATGCAGCAAGCGAGT |
| SorEd-63 | CGGGTCTCACTTCGGTGCTGTCCGGGCCCATAAGTTTTAGAGCTAGAAATAGCAAGTT |
| SorEd-64 | TACCGCTACGGTGCCACCGTACGGATCATCTGCACAACTCTTTTAAATCAGC |

**Table S2**. Comparison between phenotypic and genotypic scores assigned to transgenic plants.

| **Plant ID.** | **Albino phenotype (%)** | **Knockout score (%)** |
| --- | --- | --- |
| pGL220-18 | 0 | 0 |
| pGL220-19 | 0 | 0 |
| pGL220-9 | 0 | 18 |
| pGL220-10 | 1 | 25 |
| pGL220-11 | 1 | 32 |
| pGL220-28 | 30 | 32 |
| pGL220-34 | 35 | 90 |
| pGL220-22 | 70 | 91 |
| pGL220-24 | 75 | 97 |
| pGL220-15 | 90 | 89 |
| pGL220-14 | 90 | 87 |
| pGL220-2 | 100 | 42 |
| pGL220-20 | 100 | 46 |
| pGL220-8 | 100 | 90 |
| pGL220-27 | 100 | 91 |
| pGL220-29 | 100 | 92 |
| pGL220-26 | 100 | 94 |
| pGL220-5 | 100 | 95 |
| pGL220-7 | 100 | 95 |
| pGL220-25 | 100 | 98 |
